# Label-free biochemical imaging and timepoint analysis of neural organoids via deep learning-enhanced Raman microspectroscopy

**DOI:** 10.1101/2025.09.29.679057

**Authors:** Dimitar Georgiev, Ruoxiao Xie, Daniel Reumann, Xiaoyu Zhao, Álvaro Fernández-Galiana, Mauricio Barahona, Molly M. Stevens

**Author notes:** Correspondance: Mauricio Barahona,; Molly M. Stevens. D.G. and R.X. contributed equally to this work. Department of Materials, Design and Manufacturing Engineering, School of Engineering, University of Liverpool, Liverpool, L69 3GH, UK.

## Abstract

Three-dimensional organoids have emerged as powerful models for studying human development, disease and drug response *in vitro*. Yet, their analysis remains constrained by standard imaging and characterisation techniques, which are invasive, require exogenous labelling and offer limited multiplexing. Here, we present a non-invasive, label-free imaging platform that integrates Raman microspectroscopy with deep learning-based hyperspectral unmixing for unsupervised, spatially resolved biochemical analysis of neural organoids. Our approach enables high-resolution mapping of cellular and subcellular structures in both cryosectioned and intact organoids, achieving improved imaging accuracy and robustness compared to conventional methods for hyperspectral analysis. Using our platform, we demonstrate volumetric imaging of a neural rosette within a neural organoid, and interrogate changes in biochemical composition during early developmental stages in intact neural organoids, revealing spatiotemporal variations in lipids, proteins and nucleic acids. This work establishes a versatile framework for high-content, label-free (bio)chemical phenotyping with broad applications in organoid research and beyond.

---

The ability to image and accurately characterise the architecture and molecular composition of cells and tissues has been central to advancing biological and medical sciences. To this end, a broad array of techniques – ranging from histological staining and immunofluorescence to omics-based profiling – has been developed, each offering distinct insights across spatial and molecular scales ^1–6^. Nonetheless, a fundamental challenge persists: most available methods are inherently invasive and/or require molecular labelling.

These constraints become particularly limiting as scientific research shifts toward complex, three-dimensional (3D) tissue models, such as organoids. Over the past decade, neural organoids derived from pluripotent stem cells have emerged as powerful biological models of the developing human brain, recapitulating key features of early developmental dynamics and organisation ^7–10^. Compared to two-dimensional (2D) neural cell culture models, neural organoids exhibit enhanced complexity and three-dimensional structures, providing a more advanced representation of tissue architecture ^7,10^, cellular diversity ^11,12^ and functional network formation ^13^. Consequently, neural organoids have spurred new opportunities for studying human brain development, disease and drug response *in vitro* ^14–17^. Yet, despite growing interest, organoid research remains constrained by the limitations of conventional methods for tissue imaging and characterisation. There is a pressing need for techniques that can interrogate spatial organisation and dynamics in intact organoid systems under physiologically relevant settings, without prior knowledge of specific molecular targets or the introduction of exogenous labels.

Currently, immunofluorescence staining combined with fluorescence microscopy is widely used as a gold standard, reference technique for visualising the morphological organisation of 2D and 3D cellular models, owing to its high specificity and spatial resolution ^18,19^. In this approach, fluorophore-conjugated antibodies are used to selectively bind to and visualise specific target molecules ^20^. Despite its strengths, however, this technique has several inherent limitations. First, while protein labelling is generally well-established, detecting other biomolecular classes, such as carbohydrates, lipids and various small molecules (e.g. drugs and metabolites), often requires more specialised probes or labelling strategies, or is not possible altogether. Additionally, immunofluorescence staining involves multiple washing and permeabilisation steps, which can disrupt cellular architecture and compromise membrane integrity, potentially leading to artefactual results ^21^. These steps may also displace small molecules such as drugs, metabolites and other transient compounds, obscuring their true spatial localisation *in situ* within cells and tissues. While other labelling strategies, such as fluorescent protein tagging, genetic modifications and dye-based staining, may help avoid these washing steps, they remain label-based (i.e. targets must be known and available), and issues with labelling non-proteins persist. Furthermore, regardless of the labelling strategy, the intrinsic spectral overlap of fluorescent markers constrains multiplexing, typically to four targets in standard fluorescence microscopy ^22^, or seven in more advanced techniques ^23,24^. Moreover, genetic modifications used in live-cell imaging to express fluorescent protein reporters can be costly, time-consuming, and may induce poorly understood side effects, such as protein aggregation or disruption of ion transport dynamics. Enabling noninvasive, label-free analysis would thus be essential to expand the potential of neural organoids in neurodevelopmental research, disease modelling, drug screening and beyond.

Raman spectroscopy (RS), a technique within vibrational spectroscopy, offers a promising alternative for biochemical analysis. Unlike fluorescence-based imaging, RS provides molecular contrast through the analysis of the intrinsic inelastic light scattering of molecules, thereby enabling non-invasive, label-free chemical characterisation with minimal sample preparation ^25–27^. Consequently, RS has become a ubiquitous analytical technique across the life and physical sciences ^28–38^, with increasing use in analysing biological specimens such as cells and tissues ^39–44^. In the context of neurodevelopment, RS has been applied to profile neural cells at various stages of differentiation. Yet, most studies to date have focused on 2D cultures of individual cell types with RS measurements acquired at a single-cell level – e.g. distinguishing neural stem cells from neurons by identifying distinct cellspecific spectral signatures ^45–49^. The application of RS to complex three-dimensional cultured tissues containing multiple cell types, such as neural organoids, remains challenging due to their high biochemical heterogeneity, which hinders signal unmixing and interpretation. Recently, Bruno et al. explored the use of supervised machine learning to classify RS spectra from cortical organoids at four predefined maturation time points (weeks 6, 12, 16 and 20) ^50^. However, their approach focused on single-point prediction and did not explore the threedimensional architecture, cellular interactions and functional complexity intrinsic to neurodevelopment. So far, the use of RS for spatially resolved analysis of neural organoid structure and maturation remains unexplored.

Our lab recently reported a technique based on RS for 3D biochemical imaging and drug distribution analysis in liver organoids ^51^. The proposed methodology relied on the acquisition of ‘tissue phantoms’ – i.e. calibrated reference spectra from biomolecules of interest, which were used to guide the estimation of chemical concentration maps across the scanned area using ordinary least squares with non-negativity constraint ^52^. This enabled the detection and visualisation of key biomolecules and drugs *in situ* in whole liver organoids. Nonetheless, the approach has two limitations. First, it assumes that target biomolecules are known, thereby restricting potential applications and undermining the label-free capabilities of RS. Second, the performance and robustness of ordinary least squares for chemical estimation can be compromised by complex mixtures, experimental noise and artefacts, and variations introduced during data calibration and preprocessing, leading to inaccurate qualitative and quantitative analysis. To address these limitations, we recently introduced a deep learning framework for RS chemometrics, using hyperspectral unmixing autoencoder neural networks, which enabled fully unsupervised, label-free (bio)chemical analysis with enhanced accuracy compared to established chemometric techniques ^53^.

Building on this work, we present here a pipeline for non-invasive, label-free biochemical imaging of both cryosectioned and intact neural organoids based on spontaneous Raman microspectroscopy and unsupervised autoencoder-based hyperspectral unmixing analysis. We show that our approach achieves improved organoid imaging compared to standard methods for RS analysis, as validated against fluorescence microscopy, and enables detailed morphochemical imaging and spatially resolved analysis in 2D and 3D with subcellular resolution. Using our platform, we demonstrate volumetric *in situ* imaging of a neural rosette structure within a neural organoid – a key morphological feature of the developing nervous system – and interrogate changes in biochemical composition during early developmental stages in intact neural organoids, revealing spatiotemporal variations in lipids, proteins and nucleic acids.

## Results

### Deep learning-enhanced Raman microspectroscopy

We begin by discussing the technical methodology behind our organoid imaging pipeline based on Raman microspectroscopy combined with autoencoder-based hyperspectral unmixing. Figure 1A provides an overview of our approach.

**Figure 1:**
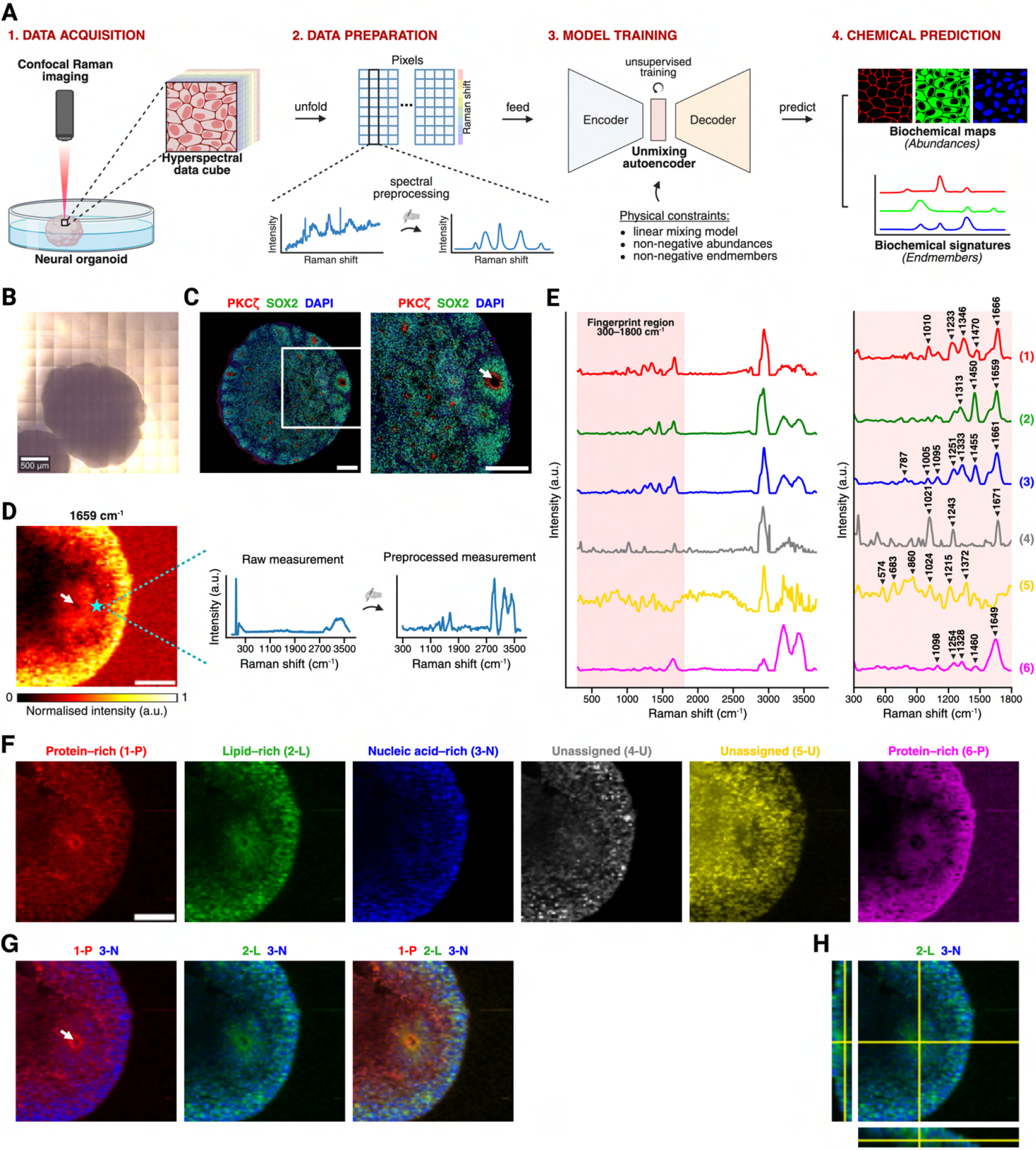
Deep learning-enhanced Raman microspectroscopy enables label-free *in situ* biochemical imaging of intact neural organoids in 2D and 3D. **A**, Schematic diagram of our organoid imaging pipeline combining Raman microspectroscopy and deep learning-enhanced hyperspectral analysis. **B**, Brightfield image of a representative day-20 neural organoid sample. Scale bar: 500 µm. **C**, Fluorescence microscopy image of a representative day-20 neural organoid sample – full organoid sample (*left*), and a zoomed-in region (*right*), with staining markers for apical domains (PKC*ζ*), neural progenitor cells (SOX2) and nuclei (DAPI). Arrow indicates the lumen of a present neural rosette. Scale bar: 200 µm. **D**, Snapshot of collected volumetric Raman imaging data from an intact day-20 neural organoid, showing the intensity profile at the 1659 cm^−1^ band (proteins and lipids) of a selected *z*-layer (*left*), and a representative tissue spectrum before and after preprocessing (*right*). Arrow indicates the lumen of a present neural rosette. Scale bar: 100 µm. **E–H**, Results of hyperspectral unmixing analysis. **E**, Derived tissue-specific endmember signatures – whole wavenumber range (*left*) and fingerprint region only (*right*). Endmembers scaled for visualisation purposes (maximum intensity set to 1). **F–G**, Corresponding abundance maps for the same *z*-layer as in **D**, including single-channel images (**F**) and representative multi-channel images (**G**). Arrow indicates the lumen of the neural rosette shown in **G**. Scale bar: 100 µm. **H**, Volumetric abundance reconstruction of two selected endmembers linked to lipid- and nucleic acid-rich species.

Raman measurements were collected from fixed neural organoid samples by performing point-wise imaging scans across predefined regions of interest. Volumetric scans were generated by acquiring sequential scans along the axis perpendicular to the imaging plane (e.g. sampling at different depths along the *z*-axis). Acquired data were preprocessed to remove non-specific signal contributions and spectral artefacts, such as cosmic spikes, autofluorescence baselines and dark noise (see Methods and Materials). This was done using RamanSPy – an open-source Python toolbox for RS data analysis we recently developed ^54^.

After preprocessing, hyperspectral unmixing was performed to derive constituent spectral components (endmembers) and quantify their relative contributions (abundances) in each measurement ^55,56^. This was achieved using an autoencoder neural network model designed for hyperspectral unmixing, following our previously reported approach ^53^. Briefly, the autoencoder architecture included: (i) an encoder responsible for abundance estimation, which comprised a multi-branch spectral convolutional block followed by a five-layer fully connected network, and (ii) a one-layer decoder responsible for endmember identification. Our model incorporated relevant physical constraints in the representation learning process, namely, constraining the learnt endmembers and abundances to be non-negative through the use of appropriate activation functions and weight clipping, and constraining the mixing process to linear mixing through the use of a single-layer linear decoder. Models were trained on individual scans or datasets in an unsupervised manner by minimising a reconstruction loss based on the weighted sum of the mean squared error and spectral angle divergence ^57^ between input spectra and output reconstructions. The number of endmembers to extract for each scan/dataset was determined using principal component analysis (PCA) ^58,59^(see Methods and Materials).

After model training, learnt endmember signatures and their corresponding abundances were derived, generating multichannel abundance images/volumes with channels rendering chemical maps of the distribution of learnt endmembers. The obtained endmembers and abundances were examined to filter out components linked to non-tissue signals, such as background and noise. The remaining tissue-specific endmembers were characterised via peak assignment, whereby prominent peaks were linked to chemical bonds to identify essential biomolecular species, such as nucleic acids, proteins, lipids and carbohydrates. Peak identification was guided by established reference materials ^40^. The corresponding abundance maps were visualised as morphochemical images showing the spatial distribution of the obtained chemical components to study organoids’ morphology and biochemical composition.

Further technical details regarding data acquisition, preprocessing and unmixing analysis are presented in Methods and Materials.

### Label-free imaging of intact neural organoids via deep learning-enhanced Raman microspectroscopy

We investigated the capabilities of our platform using Raman imaging data from neural organoids cultured in-house following an unguided differentiation protocol ^7^ (Figure S1 and Methods and Materials). Brightfield microscopy revealed spherical neural organoid samples reaching about 1–2 mm in size after 20 days in culture (Figure 1B). Furthermore, fluorescence microscopy showed characteristic morphology with neural rosettes readily visible on day 20 (Figure 1C).

As a demonstration of our Raman imaging platform, we performed volumetric Raman imaging of a tissue region within an intact day-20 neural organoid. Specifically, a *z*-stack was acquired with an axial step size of 10 µm in *z* (organoid depth), with each layer representing an 80 × 80 imaging scan with a lateral pixel size of 5 µm in the *x*-*y* plane. Five consecutive layers along the *z*-axis were used for unmixing analysis, covering tissue areas up to approximately 70 µm deep into the sample. Figure 1D shows the intensity profile at 1659 cm^−1^ of one of the five layers after preprocessing, a band associated with proteins (amide I) and lipids (C=C stretching). Although the overall outline of the organoid was visible, finer structures were difficult to resolve using univariate analysis of single Raman bands.

To address this issue, we performed multivariate hyperspectral unmixing using our deep learning-enhanced Raman microspectroscopy pipeline. Our analysis yielded six tissuespecific spectral endmembers, which we characterised via peak assignment into major biomolecular classes based on characteristic Raman vibrations in the fingerprint region 300 cm^−1^ to 1800 cm^−1^ (Figure 1E; extended data in Figure S2). The corresponding abundance maps were visualised as morphochemical images of the scanned organoid sample (Figure 1F–G). Endmember 1 (red) exhibited a spectral signature characteristic of protein species, with prominent peaks at 1010 cm^−1^ (phenylalanine), 1233 cm^−1^ (amide III), 1346 cm^−1^ (CH deformation), 1470 cm^−1^ (CH_2_ bending, C=N stretching) and 1666 cm^−1^ (amide I). The abundance map showed broad distribution throughout the organoid tissue, likely corresponding to cytoplasmic regions. Slightly increased intensity was observed around a structure resembling the apical domain of a neural rosette, possibly indicating membrane-associated protein structures. Endmember 2 (green) was characterised by strong lipid-associated peaks at 1313 cm^−1^ (CH_2_/CH_3_ twisting), 1450 cm^−1^ (CH_2_ bending) and 1659 cm^−1^ (C=C stretching), suggesting the presence of lipid-rich compartments such as cellular membranes and lipid bodies. This component was similarly broadly distributed across the tissue, with increased signal in the outer layers of the sample, and around the apical domain of the neural rosette, forming a radially oriented pattern. Endmember 3 (blue) displayed main nucleic acid vibrations, including 787 cm^−1^ (O–P–O stretching) and 1095 cm^−1^ (PO_2_^−^ groups), alongside protein-related peaks at 1005 cm^−1^, 1251 cm^−1^, 1455 cm^−1^ and 1661 cm^−1^. This is consistent with cell nuclear regions, which is further supported by visual inspection when overlaid with the abundance maps of endmembers 1 and 2. Endmembers 4 (grey) and 5 (yellow) presented less conventional profiles, suggesting more complex biomolecular mixtures. Endmember 4 included peaks at 1021 cm^−1^ (glycogen), 1243 cm^−1^ (proteins) and 1671 cm^−1^ (proteins/lipids), and its spatial distribution overlapped with that of endmember 3. Endmember 5 displayed a noisier spectrum, with higher abundance within the interior of the organoid. Prominent peaks were associated to proteins (574 cm^−1^, tryptophan/cytosine; 860 cm^−1^, C–C stretching; 1215 cm^−1^, amide III) and carbohydrates (1024 cm^−1^, glycogen; 1372 cm^−1^, CH deformation). Lastly, endmember 6 (magenta) showed a broadly distributed component, with bands related to proteins (1254 cm^−1^, 1328 cm^−1^, 1460 cm^−1^ and 1649 cm^−1^) and nucleic acids (1098 cm^−1^).

Beyond 2D imaging, our approach also enables biochemical analysis in three dimensions. To illustrate this, we generated a volumetric rendering of the imaged organoid by overlaying the two derived components related to lipid-rich (endmember 2) and nucleic acid-rich (endmember 3) species as markers of nuclear and membrane regions, respectively. The reconstruction, displayed in Figure 1H, allowed the visualisation and analysis of the three-dimensional organisation and architecture of the imaged organoid tissue, including that of the present neural rosette (rosette lumen indicated with an arrow in Figure 1D). The derived organoid image using RS showed characteristic morphology, correlated with fluorescence microscopy (Figure 1C), yet was achieved without exogenous labelling or tissue sectioning. This demonstrates that our Raman-based imaging platform can effectively delineate organoid composition *in situ* and generate 2D and 3D morphochemical maps of key biochemical species in intact neural organoid samples in a fully label-free manner. Building on this proof of concept, we proceed to higher-resolution Raman imaging in the following sections.

### Experimental validation and optimisation against fluorescence microscopy

To further validate that our method produces accurate morphochemical organoid reconstructions, we evaluated its performance against fluorescence microscopy and standard methods for hyperspectral RS analysis. To this end, defined tissue regions within two day-30 neural organoid cryosections were imaged using confocal Raman microspectroscopy, followed by immunostaining and fluorescence microscopy (Figure 2A). Raman imaging was performed first to avoid potential signal interference from staining molecules. To ensure morphological diversity, we selected areas containing both distinct neural rosette structures and surrounding neural tissue for downstream imaging (Figures 2B, S3A and S5A). Raman imaging was performed over areas of 600 µm × 600 µm and 400 µm × 400 µm from the two respective sections, with a pixel size of 2 µm (Figures S3C and S5C). Subsequently, the organoid sections were stained for nuclei (DAPI), cell membranes (DiO), neural progenitor cells (SOX2) and neurons (TUJ1), and fluorescence microscopy images were taken to validate and correlate with organoid morphology predicted by RS (Figures S5B and S3B).

**Figure 2:**
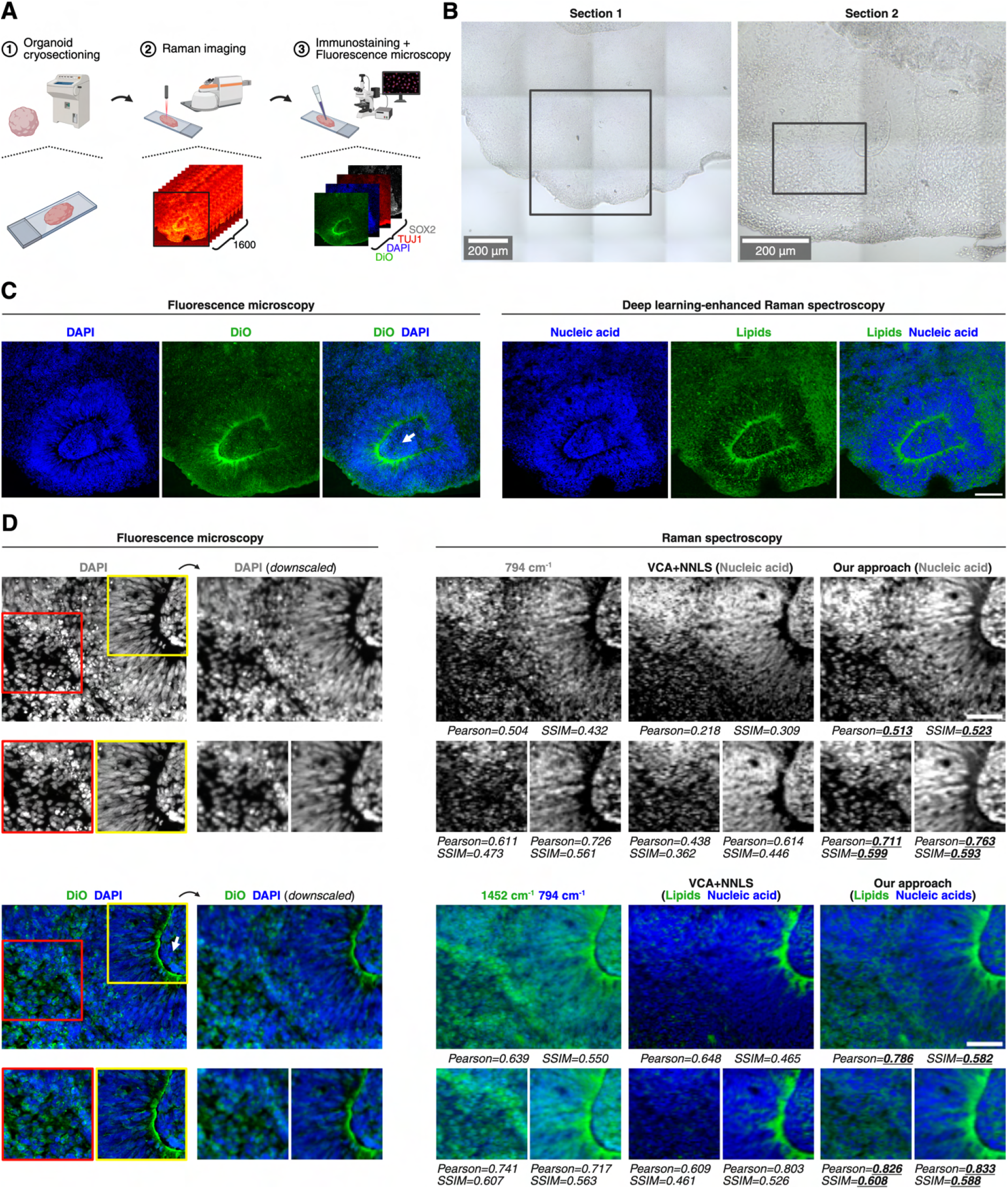
Deep learning-enhanced Raman microspectroscopy enables accurate morphochemical imaging of neural organoids as validated against fluorescence microscopy. **A**, Schematic diagram of our validation strategy, which involved the collection of paired Raman imaging and fluorescence microscopy data from neural organoid cryosections. **B**, Brightfield microscopy images of the two day-30 neural organoid cryosections imaged for validation. Marked areas show the regions used for validation in **C** and **D**, respectively. Scale bars: 200 µm. **C**, Imaging results on first organoid section. Comparison of fluorescence microscopy images of DAPI (nuclei) and DiO (membranes) (*left*), and RS-based abundance reconstructions of nucleic acid and lipids derived with our autoencoder pipeline (*right*). Arrow indicates the lumen of the present neural rosette. Scale bar: 100 µm. **D**, Imaging results on second organoid section. Comparison of fluorescence microscopy images of DAPI (nuclei) and DiO (membranes) (*left*), and RS-based abundance reconstructions of nucleic acid and lipids derived with three methods for Raman imaging, including univariate analysis, unmixing analysis with VCA and NNLS, and our autoencoder pipeline. Similarity metrics, namely Pearson’s correlation coefficient (Pearson) and structural similarity index measure (SSIM), were performed with respect to the respective downscaled fluorescence images. The highest achieved similarity scores are indicated by underlined bold text. Arrow indicates the lumen of the present neural rosette. Scale bar: 50 µm.

Using our autoencoder-based pipeline, we analysed the acquired RS data from the first organoid section to infer its biochemical composition. Our method extracted four tissuespecific endmembers, which we associated with contributions from lipids, nucleic acids and proteins (Figures S3D–E and S4). Figure 2C shows a side-by-side comparison between the predicted abundance maps of the two endmembers related to nucleic acid-rich and lipid-rich species and the corresponding fluorescence microscopy images of nuclei (DAPI) and cell membranes (DiO), respectively. We observed that our RS imaging pipeline accurately reconstructed individual nuclei, aligning with the DAPI staining across the imaged tissue (Figures 2C (right) and S3F). It also effectively captured important membrane features, including both fine cellular membranes and prominent structures such as the apical domain of the visible neural rosette. Qualitatively, we observed strong correspondence between the lipid-rich component obtained with RS and the DiO signal around the apical domain of the rosette and with the TUJ1 signal outside the rosette (Figures S3B and S3E).

Having shown that our method accurately reconstructs key cellular components in cryosectioned neural organoid samples, we used the data collected from the second organoid section to perform a series of comparative analyses to test different versions of our autoencoder-based pipeline, including using different training losses (Figure S8) and endmember initialisation procedures (Figure S9). Following this, we compared the unmixing performance of our developed approach against a range of standard chemometric methods for multivariate analysis. This included methods for dimensionality reduction (PCA ^58,59^, non-negative matrix factorisation (NMF) ^60^), clustering (k-means clustering ^61,62^), and hyperspectral unmixing (N-FINDR ^63^ and vertex component analysis (VCA) ^64^ for endmember identification, applied together with non-negative least squares (NNLS) ^52^ for abundance estimation).

Our autoencoder-based pipeline derived six tissue-specific endmembers, which we characterised as lipid-, nucleic acid- and protein-rich species (Figures S5D and S6). Visual examination of the estimated abundance maps revealed major cellular and subcellular features, including membranes, nuclei, cytoplasmic regions and apical domains of neural rosettes (Figure S5E). Compared to standard methods, our approach offered improved robustness to artefacts and noise and produced more discriminative and interpretable abundance maps, enabling better-delineated imaging reconstruction of organoid morphology (Figure S7).

To quantitatively assess the performance of our autoencoder-based pipeline, we evaluated the image similarity between Raman-based reconstructions and fluorescence microscopy images in an overlapping organoid region (Figure 2D). To this end, we spatially aligned both modalities and computed the Pearson’s correlation coefficient (Pearson) and structural similarity index measure (SSIM) ^65^ between RS-derived abundance maps of nuclei and membranes and fluorescence images of DAPI and DiO, respectively. We benchmarked the performance of our method against two conventional chemometric approaches: (i) univariate analysis, and (ii) multivariate hyperspectral unmixing using VCA and NNLS. Univariate analysis was performed using the intensity profiles at 794 cm^−1^ and 1454 cm^−1^, which we found to be indicative of nuclear and lipid-rich membrane regions, respectively (Figure S5D).

Univariate analysis using the 794 cm^−1^ band yielded nuclear reconstructions which were generally aligned with the DAPI signal. However, visual inspection revealed irregular structures, which impeded the precise localisation of individual nuclei (Figure 2D, right, first column). Major membrane structures, such as the apical domain of the imaged rosette, were also recovered via the 1454 cm^−1^ band, yet finer membrane features were not visually discernible, obscured by high background and non-specific signal. This could be attributed to the overlap with vibrations of proteins in the same spectral range or non-specific intensity variations due to acquisition or preprocessing settings, underscoring the need for more robust multivariate chemometric approaches ^66^. Multivariate unmixing analysis with VCA and NNLS showed low reconstruction accuracy, with further reduced Pearson and SSIM scores compared to univariate analysis for both nuclei and membranes (Figure 2D, right, second column). Visually, nuclear reconstructions were similarly non-distinct, with poorly defined nuclei. Moreover, while major membrane structures like the apical domain of the present rosette were recovered with higher specificity, finer membrane features were consistently undetected. By contrast, as shown in Figure 2D (right, third column), our autoencoder-based pipeline achieved the highest reconstruction accuracy across all evaluations – both quantitatively, with superior Pearson and SSIM scores, and qualitatively, with more interpretable and biologically relevant output. Nuclear regions were recovered with higher accuracy, with visually better-defined nuclei and lower non-specific signal. Membrane-associated structures were also more clearly delineated, enabling the visualisation of both large-scale features such as apical domains and individual cellular membranes.

Furthermore, our approach demonstrated improved robustness to experimental noise, maintaining consistent reconstruction quality under reduced denoising conditions. In comparison, the performance of conventional methods deteriorated markedly, yielding noisier, less interpretable spectral components and abundance profiles, or failing to converge to a solution during unmixing altogether (Figure S10).

### Biochemical imaging of intact neural organoids at subcellular resolution

After demonstrating that our method achieves accurate subcellular imaging of 2D cryosectioned neural organoid tissue, we performed higher-resolution Raman imaging of a tissue region within an intact 3D neural organoid. The ability to resolve and visualise subcellular structures in intact organoid tissues in their native environment is critical for interrogating important neurodevelopmental processes, such as nuclear positioning, polarity establishment and lumen formation, and studying their effect on brain tissue organisation, architecture and function.

To this end, we acquired a larger 400 × 346 Raman imaging scan taken with 1 µm pixel size from an intact day-10 neural organoid sample (Figure 3A). Using our Raman imaging pipeline, we analysed the data and identified six tissuespecific components. The spectral endmembers and corresponding abundance maps are presented in Figure 3B and Figure 3C, respectively (extended data in Figure S11). Endmember 1 (red) exhibited a lipid-rich profile with prominent peaks at 1077 cm^−1^ (C–C and C–O stretching, phospholipids), 1305 cm^−1^, 1448 cm^−1^ and 1657 cm^−1^. Its abundance map outlined membrane-like structures, along with more confined bright spots resembling lipid droplets. Endmember 2 (grey) was characterised by vibrations associated with both protein and lipid species, with peaks at 1000 cm^−1^, 1124 cm^−1^ (C– C and C–N stretching), 1249 cm^−1^, 1311 cm^−1^, 1454 cm^−1^ and 1654 cm^−1^. Its abundance was observed in regions partially overlapping with membranes, as well as in isolated domains, possibly indicating cytoplasmic or extracellular matrix compartments. Endmember 3 (blue) was marked by strong nucleic acid bands at 790 cm^−1^ and 1095 cm^−1^, alongside protein-associated bands at 1005 cm^−1^, 1249 cm^−1^, 1328 cm^−1^, 1455 cm^−1^ and 1659 cm^−1^. As in previous analyses, the abundance map revealed well-defined spherical compartments consistent with individual cell nuclei. Endmember 4 (green) presented a distinct protein-rich signature with peaks at 1005 cm^−1^, 1251 cm^−1^, 1454 cm^−1^ and 1662 cm^−1^, along with vibrations associated with ring breathing modes in nucleic acid bases at 1573 cm^−1^. The abundance map highlighted cellular-level compartments resembling cytoplasmic regions, as well as small punctate structures within nuclei with high signal intensity, potentially corresponding to biomolecular condensates. Endmember 5 (yellow) displayed a mixed profile comprising bands linked to proteins (1259 cm^−1^, 1326 cm^−1^, 1453 cm^−1^ and 1662 cm^−1^) and nucleic acid (1095 cm^−1^). This component appeared more diffusely distributed, with substantial spatial overlap with endmember 3. Endmember 6 (purple) presented a noisier signature, with carbohydrate-associated bands at 493 cm^−1^ (glycogen), 850 cm^−1^ (C–O–C skeletal mode, glycogen) and 938 cm^−1^ (C–C stretching), along with peaks linked to proteins and nucleic acids at 754 cm^−1^ and 1103 cm^−1^. The abundance overlapped with endmember 4, particularly around nuclear condensates, potentially reflecting glycoproteins within cell nuclei.

**Figure 3:**
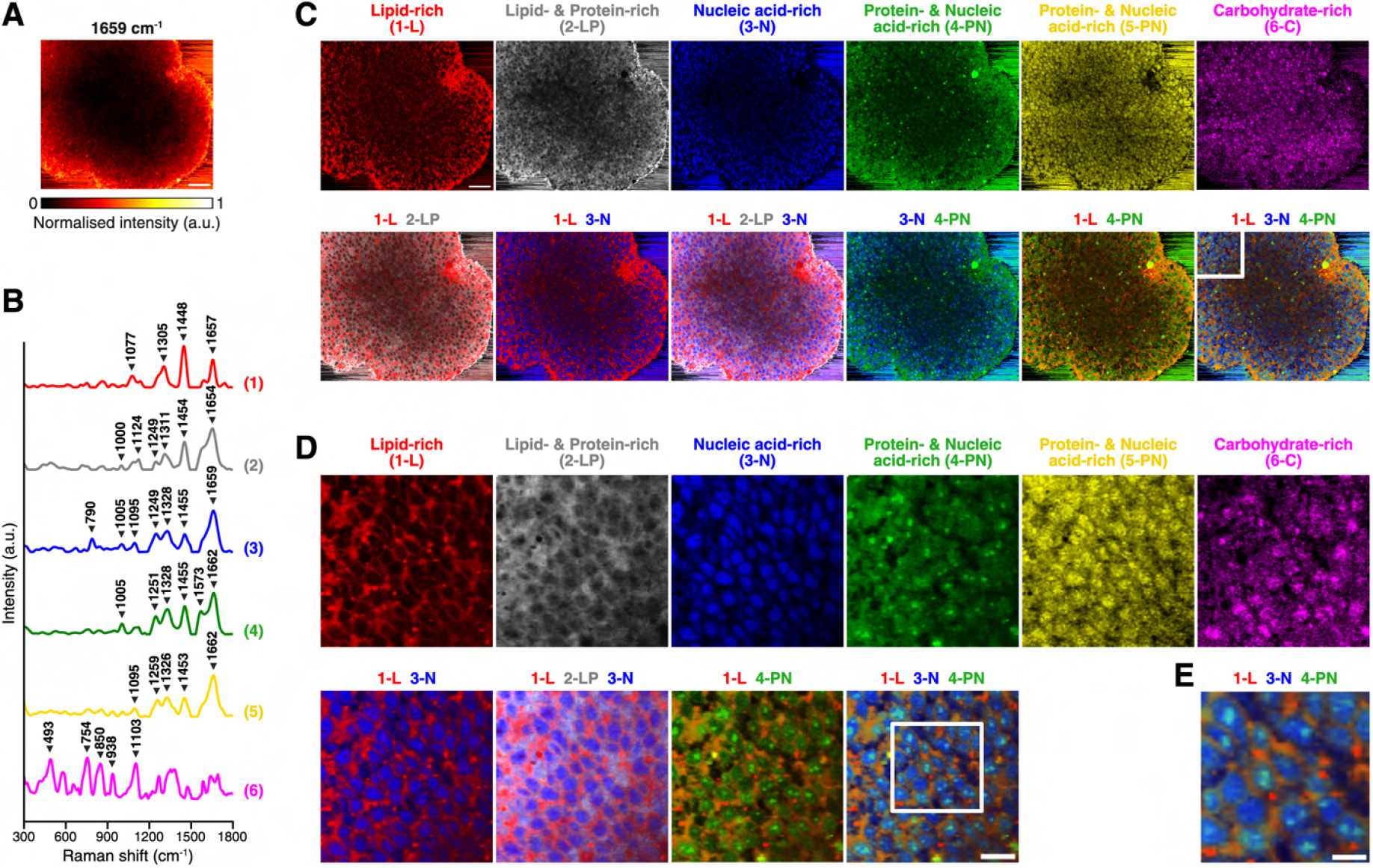
Biochemical imaging of intact neural organoids at subcellular resolution. **A**, Snapshot of collected Raman imaging data from an intact day-10 neural organoid sample, showing the intensity profile at the 1659 cm^−1^ band (proteins and lipids). Scale bar: 50 µm. **B–E**, Results of hyperspectral unmixing analysis. **B**, Derived tissue-specific endmember signatures. Endmembers scaled for visualisation purposes (maximum intensity set to 1). **C**, Corresponding abundance maps. Scale bar: 50 µm. **D**, Zoomed-in abundance images of a selected area (area marked in **C**, bottom right panel). Scale bar: 20 µm. **E**, Further zoomed-in abundance image (area marked in **D**). Scale bar: 10 µm.

To further inspect the spatial reconstructions produced by our imaging pipeline, we examined magnified fields from selected regions within the imaged organoid tissue (Figures 3D– E). The resulting reconstructions revealed consistent, welldefined cellular features, which delineated individual cells and subcellular structures resembling nuclei, cytoplasmic regions, cellular membranes and nuclear condensates.

These results demonstrate the effectiveness of our imaging platform across scales and resolutions, allowing us to resolve complex biochemical heterogeneity and interrogate key morphological features at cellular and subcellular scales in a single, label-free imaging modality.

### Volumetric *in situ* imaging of a neural rosette within an intact neural organoid

Next, we extended our subcellular imaging analysis to 3D and used our pipeline to perform volumetric *in situ* imaging of a neural rosette structure within an intact neural organoid.

Neural rosettes are important organisational units during neural organoid development, which recapitulate key aspects of neural tube formation in the embryonic central nervous system ^67^. They typically form within the first two weeks of neural induction and appear as radially organised, layered structures composed primarily of neuroepithelial cells at early stages. As development progresses, these progenitor cells give rise to differentiated neural lineages, including neurons and glial cells. To illustrate this, we performed fluorescence microscopy of a day-20 neural organoid tissue section, with markers for nuclei (DAPI), neurons (MAP2), neural progenitor cells (SOX2) and apical domain (PKC*ζ*). The resulting images, shown in Figure 4A, revealed multiple neural rosettes with characteristic radial structure. We observed an abundance of neural progenitor cells densely organised within the rosettes, with surrounding layers of differentiating neurons.

**Figure 4:**
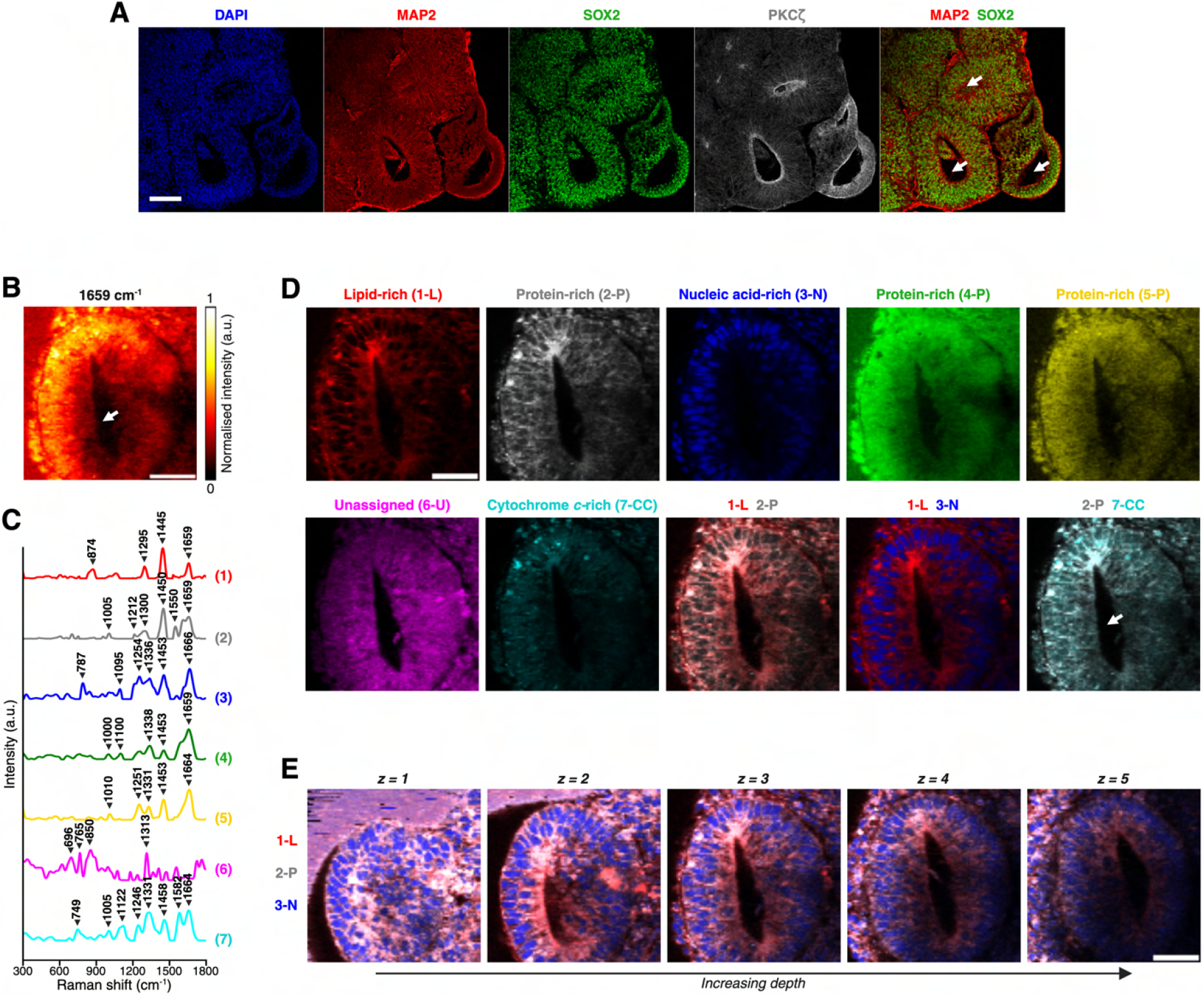
Volumetric *in situ* biochemical imaging of a neural rosette within an intact neural organoid. **A**, Fluorescence microscopy images of a day-20 neural organoid tissue section, showing characteristic neural rosette structures, with markers for nuclei (DAPI), neurons (MAP2), neural progenitor cells (SOX2) and apical domain (PKC*ζ*). Arrows indicate the lumens of developed neural rosettes. Scale bar: 100 µm. **B**, Snapshot of collected volumetric Raman imaging data from an intact day-20 neural organoid sample (selected *z*-layer), showing the intensity profile at the 1659 cm^−1^ band (proteins and lipids). Arrow indicates the lumen of the present neural rosette. Scale bar: 50 µm. **C–E**, Results of hyperspectral unmixing analysis. **C**, Derived tissue-specific endmember signatures. Endmembers scaled for visualisation purposes (maximum intensity set to 1). **D**, Corresponding abundance maps for a representative *z*-layer. Arrow indicates the lumen of the present neural rosette, as also shown in **B**. Scale bar: 50 µm. **E**, Image reconstructions of selected endmember components at different depths in *z*. Scale bar: 50 µm.

Following this, we investigated whether our Raman-based imaging platform could be used to visualise and analyse the structure of a neural rosette without the need for tissue sectioning and exogenous labelling. To this end, we acquired a volumetric RS scan of a neural rosette within an intact day-20 neural organoid, consisting of five imaging slices in *z* (organoid depth), with each slice measuring 95 × 95 pixels with a pixel size of 2 µm. Figure 4B displays the intensity profile at the 1659 cm^−1^ peak (proteins and lipids) of one of the collected layers, outlining a characteristic neural rosette structure. Our autoencoder-based unmixing analysis yielded six tissue-specific components (Figure 4C–D, extended data in Figure S12). Endmember 1 (red) presented a lipid-rich profile with prominent peaks at 874 cm^−1^ (phospholipids), 1295 cm^−1^ (CH_2_ deformation), 1445 cm^−1^ and 1659 cm^−1^. The corresponding abundance map highlighted structures resembling cellular membranes and lipid bodies. Endmember 2 (grey) contained features associated with proteins and lipids, with peaks at 1005 cm^−1^, 1300 cm^−1^, 1450 cm^−1^, 1550 cm^−1^ (C=C stretching, tryptophan) and 1659 cm^−1^. Its signal overlapped with that of endmember 1, appearing in membrane-like regions, but also extended into the cell interior, possibly indicating cytoplasmic and structural proteins. Endmember 3 (blue) showed a distinct nuclear signature, with nucleic acid-associated vibrations at 787 cm^−1^ and 1095 cm^−1^, along protein bands at 1254 cm^−1^, 1336 cm^−1^, 1453 cm^−1^ and 1666 cm^−1^. This component exhibited strong nuclear specificity, outlining well-defined spherical compartments consistent with individual cell nuclei. Endmembers 4 (green) and 5 (yellow) shared similar spectral features dominated by protein-related peaks around ∼1005 cm^−1^, ∼1100 cm^−1^, ∼1255 cm^−1^, ∼1335 cm^−1^, ∼1453 cm^−1^ and ∼1660 cm^−1^, suggesting general protein composition. Endmember 6 (magenta) displayed a less conventional, noisier profile with peaks at 696 cm^−1^ (potentially related to cholesterol), 765 cm^−1^ (pyrimidine ring breathing in nucleic acids), 850 cm^−1^ (C–O–C skeletal mode in carbohydrates, glycogen) and 1313 cm^−1^ (CH_2_/CH_3_ twisting in lipids and collagens, or guanine in nucleic acid). This component was generally broadly distributed, with some confined regions with higher abundance intensity. Lastly, endmember 7 (cyan) showed a signature well-aligned with mitochondria, featuring peaks associated with cytochrome *c* at 749 cm^−1^, 1122 cm^−1^ and 1582 cm^−1^, alongside protein-related bands at 1005 cm^−1^, 1246 cm^−1^, 1331 cm^−1^, 1458 cm^−1^ and 1664 cm^−1^. This component was observed in cytoplasmic regions, in proximity to nuclei, consistent with intracellular localisation of mitochondria.

To examine the three-dimensional structure of the scanned rosette, we overlaid the abundance maps of the derived lipid-rich (endmember 1), protein-rich (endmember 2) and nucleic acid-rich (endmember 3) components across all five *z*-layers. The resulting images, presented in Figure 4E, demonstrated that our imaging pipeline can effectively reconstruct key subcellular features – such as individual cellular membranes and nuclei – across all five imaging scans, enabling the interrogation of cellular architecture and organisation within the intact neural rosette in both 2D and 3D.

### Tracing changes in biochemical composition during early neurodevelopment in intact neural organoids

During maturation, neural organoids recapitulate key stages of early human brain development, including the formation of distinct progenitor zones and the emergence of layered neural architectures ^11,12,68,69^. To better understand the biochemical dynamics accompanying tissue differentiation and organisation, we applied our imaging pipeline to trace changes in organoid composition during neural organoid development.

To this end, we acquired Raman imaging data from organoid samples at five developmental time points – days 5, 10, 20, 30 and 40 (Figure 5A). In total, we collected 72 Raman imaging scans from organoids grown in three independent culture batches, with scan numbers ranging between 11 and 17 per time point (Figure S14A). Each scan represented a 20 ×160 image across the outermost tissue layers, which normally contain the most metabolically active and viable cells ^70,71^. Measurements were acquired with a lateral pixel size of 5 µm in the *x*-*y* plane and an axial step size of 10 µm along the *z*-axis, thereby imaging up to 800 µm across and 200 µm deep into the organoid samples. Scans were preprocessed and hyperspectral unmixing analysis was performed, yielding eight tissue-specific endmember species (Figure 5B; extended data in Figure S13). The corresponding abundance profiles were analysed to examine relative biochemical trends across developmental groups. Figure 5C displays the mean abundance of each component across time points, after background removal and area normalisation (Figure S14B).

**Figure 5:**
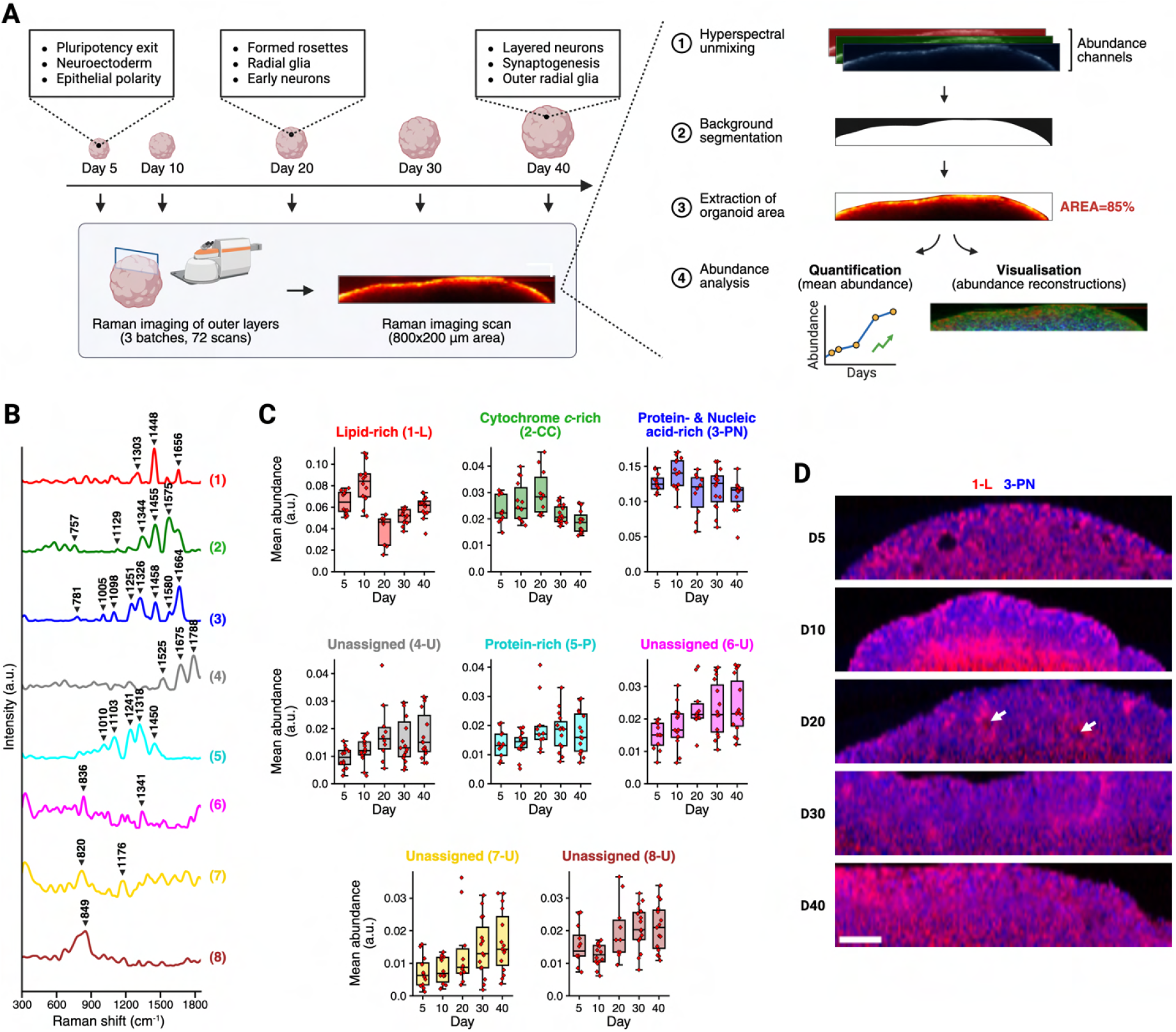
Tracing changes in biochemical composition during early neurodevelopment in intact neural organoids. **A**, Schematic diagram of our timepoint study, depicting data acquisition (*left*) and analysis (*right*). **B–D**, Results of hyperspectral unmixing analysis. **B**, Derived tissue-specific endmember signatures. Endmembers scaled for visualisation purposes (maximum intensity set to 1). **C**, Quantification of the corresponding abundances over time. Sample sizes: day 5 (*n* = 12), day 10 (*n* = 14), day 20 (*n* = 10), day 30 (*n* = 15), day 40 (*n* = 15) from *N* = 3 independent organoid culture batches. **D**, Abundance reconstructions of selected organoid samples across the five timepoint groups. Arrows indicate structures resembling developed neural rosettes. Scale bars: 100 µm

Endmember 1 (red) exhibited a distinct lipid-rich spectral profile, characterised by prominent peaks at 1303 cm^−1^, 1448 cm^−1^ and 1656 cm^−1^, and appeared broadly distributed with high abundance across all organoids and time points (Figure S15A). Still, distinct spatial and temporal variations in its distribution across time points were observed. On day 5, the signal appeared relatively homogeneous, with slightly elevated intensity in small compartments near the outer layers. By day 10, its intensity increased markedly, particularly around the inner layers and surface. The high abundance in early time points may reflect shifts in lipid metabolism during early neurogenesis, in line with previous findings that lipid accumulation is essential for neural stem/progenitor cell activity and proliferation ^72^. Following this, a sharp decline in signal was observed in day 20 organoids, where discrete, localised regions of elevated intensity emerged, likely corresponding to developed neural rosette structures. Between days 30 and 40, the signal progressively recovered, once again displaying a more uniform and widespread distribution. This gradual recovery may be linked to the expansion and functional maturation of neuronal populations, which rely on lipids not only as essential structural components of membranes but also as signalling molecules with key regulatory functions ^73,74^. This is further supported by our observation that lipid-rich endmembers are well co-localised with TUJ1-positive regions – a marker of neurons (Figure S3). In contrast, endmember 2 (green) presented a mixed spectral profile, with peaks linked to cytochrome *c* (757 cm^−1^, 1129 cm^−1^, 1575 cm^−1^) and proteins (1344 cm^−1^, 1455 cm^−1^). This signature suggested regions enriched in mitochondria, and showed an abundance profile with an inverted U-shaped trajectory, gradually increasing from day 5 to day 20, followed by a decline at later time points. Despite these temporal changes, the spatial distribution remained generally uniform across the organoid tissue throughout the sampled time course. Endmember 3 (blue) comprised vibrational features associated with nucleic acids (781 cm^−1^, 1098 cm^−1^, 1580 cm^−1^) and proteins (1005 cm^−1^, 1251 cm^−1^, 1326 cm^−1^, 1458 cm^−1^, 1664 cm^−1^), likely corresponding to cytoplasmic and nuclear regions. Its abundance was generally stable across developmental stages, with a slight downward trend. A similar progression was reported by Bruno et al., who observed a gradual decrease in nucleic acid and protein features in maturing cortical organoids ^50^. Visually, this component was highly abundant across all organoid samples, with a generally uniform distribution throughout organoids, and slightly elevated intensity in the outer tissue layers (Figure S15B). Endmembers 4–8 exhibited less conventional spectral profiles, with lower relative abundance predominantly present in deeper organoid regions (Figure S13C). They shared similar developmental trajectories, each showing a gradual increase in abundance over time. While prominent peaks within these endmembers allow for tentative biochemical assignments, further validation is required to determine whether these components reflect representative biological features, or traces arising from higher noise levels in deeper tissue due to depth-dependent signal attenuation. Endmember 4 (grey) featured two bigger peaks at 1675 cm^−1^ and 1788 cm^−1^, likely attributable to protein species, and a smaller peak 1525 cm^−1^, potentially indicating contributions from carotenoids (C–C and C=C stretching). Endmember 5 (cyan) was characterised by distinct protein-related peaks at 1010 cm^−1^, 1103 cm^−1^, 1241 cm^−1^, 1318 cm^−1^ and 1450 cm^−1^.

Endmembers 6 (magenta) and 7 (yellow) exhibited noisier profiles, featuring multiple, less-defined peaks. Some of the more pronounced vibrations included peaks at 820 cm^−1^, 836 cm^−1^, 1176 cm^−1^ and 1341 cm^−1^, all related to proteins. Lastly, endmember 8 (brown) showed a single dominant peak at 849 cm^−1^, potentially corresponding to the C–O–C skeletal vibrations of saccharides. Bruno et al. reported a similar peak around 850 cm^−1^ as a prominent predictive marker of cortical organoid maturation, showing a significant, progressive increase between weeks 6 and 20^50^. The authors attributed this feature to glycogen and associated its rise with metabolic changes accompanying neuronal differentiation.

To illustrate the dynamic morphochemical composition of neural organoids during early neurodevelopment, we further generated spatial reconstructions of selected biochemical components across organoid samples at different developmental stages. Figure 5D displays reconstructions of representative samples from each of the five time points, highlighting the two selected components associated with lipids, and proteins and nucleic acids, respectively (see Figure S13C for other components). These visualisations enabled us to trace the evolving spatial organisation and architecture of neural organoids across their development, providing insights into neural organoid maturation and further supporting the quantitative trends observed in Figure 5C.

Combined, our results serve as a proof of concept of the feasibility of label-free, RS-based biomolecular tracking of spatiotemporal dynamics in intact neural organoids during development.

## Discussion

We have developed here a method for non-invasive, label-free organoid imaging based on Raman microspectroscopy combined with unsupervised autoencoder-based hyperspectral analysis. We showed that our approach is effective at discerning key biochemical components in unlabelled Raman spectroscopy measurements, enabling 2D and 3D visualisation of structural and morphological features in both cryosectioned and intact neural organoids and at different scales and resolutions. Using our imaging platform, we studied structural and biochemical features of early neurodevelopment in neural organoids and demonstrated the ability to image neural rosettes within intact organoid samples in three dimensions. Furthermore, we studied changes in the biomolecular composition of intact neural organoids during early developmental stages, observing distinct spatiotemporal dynamics as organoids mature in major biomolecular classes, including lipids, proteins and nucleic acids.

By eliminating the need for destructive preparation steps and external labels, our workflow offers simpler sample preparation that does not compromise organoid integrity, with potential for live-organoid analysis. This opens new avenues for research into early neurodevelopment, but also in developmental diseases and drug discovery, where our approach could allow studying disease aetiology in neural organoid disease models or visualising drug distribution *in situ*. We note that our pipeline can also be applied to other types of samples, including other types of organoids and tissues. Beyond imaging, it can also be used as a feature extraction technique in RS for downstream applications – e.g. organoid phenotyping, disease classification, drug response modelling.

In this study, we used spontaneous RS to validate our pipeline under stable, reproducible conditions using fixed neural organoids. Although inherently time-consuming, spontaneous RS offers high spectral resolution and chemical specificity, and is more broadly accessible than faster, more advanced RS modalities – making it well-suited for method development and benchmarking. Importantly, our unmixing analysis framework is also compatible with emerging broadband coherent Raman scattering modalities ^75^, which support video-rate imaging over wide spectral ranges. For example, Vernuccio et al. recently reported a microscopy system based on coherent anti-Stokes Raman scattering capable of full-spectrum acquisitions over large fields of view at millisecond dwell times, combined with standard methods for unmixing (N-FINDR and NNLS) ^76^. In this context, spectral unmixing remains a key step in the analysis pipeline, and future work integrating our method with more advanced RS techniques could allow fast, high-throughput analysis of biological specimens with improved chemical accuracy – a major step toward realtime, label-free imaging of live organoids.

Finally, we note that although derived endmembers were characterised using established Raman bands, some peak assignments remain partially ambiguous due to the biochemical complexity of neural organoids and depth-dependent signal attenuation. Future work integrating Raman imaging with complementary imaging modalities could further improve qualitative and quantitative interpretation. Nonetheless, our work demonstrates the feasibility of label-free, RS-based biochemical imaging and analysis in complex, three-dimensional neural organoid systems.

## Methods and Materials

### Organoid culture and preparation

Human embryonic stem cells (WA09, WiCell) were cultured feeder-free on hES-qualified Matrigel (354277, Corning) coated plates in a 5% CO_2_ incubator at 37^°^C. mTeSR1 medium (STEMCELL Technologies) was used for daily cell maintenance according to manufacturer instructions.

To generate neural organoids, cells were dissociated into single cells using Accutase (STEMCELL Technologies), resuspended in mTeSR1 supplemented with 50 µm Y-27632 (S1049, Selleck Chemicals) and 1% (v/v) Penicillin/Streptomycin (15140122, Thermo Fisher Scientific), and then added to an ultra-low-binding 96-well plate (Corning) at 9000 cells/150 µm per well. On day 3, 100 µm of the media was removed and 150 µm fresh mTeSR1 containing 1% (v/v) Penicillin/Streptomycin was added. From day 5 onwards, media was changed to neural induction (NI) media, composed of Dulbecco’s Modified Eagle Medium (DMEM)/F12 (Thermo Fisher Scientific) supplemented with 1 µm/mL heparin (Sigma-Aldrich), 1% (v/v) Penicillin/Streptomycin, 1%(v/v) GlutaMAX (Thermo Fisher Scientific), 1% (v/v) N2 supplement (Thermo Fisher Scientific), and 1% (v/v) MEM-NEAA (Thermo Fisher Scientific). On day 10, organoids were embedded in Matrigel (354234, Corning) and transferred to 10 cm petri dishes. From day 12 to day 18, the culture medium was switched to Improved-A medium consisting of a 1: 1 mixture of DMEM/F12 and Neurobasal Medium (Thermo Fisher Scientific) supplemented with 1% (v/v) Penicillin/Streptomycin, 1% (v/v) B27 supplement without vitamin A (Thermo Fisher Scientific), 1% (v/v) GlutaMAX, 0.5% (v/v) N2 supplement, 1*/*4000 (v/v) insulin (I9278, Sigma-Aldrich), 0.5% (v/v) MEM-NEAA, and 1 mg/mL sodium bicarbonate. Organoids were fed with fresh Improved-A medium every 3 days – i.e. on days 12 and 15. From day 18 onwards, the petri dishes were placed on an orbital shaker, and the culture medium was replaced with Improved+A medium consisting of a 1: 1 mixture of DMEM/F12 and Neurobasal Medium (Thermo Fisher Scientific) supplemented with 1% (v/v) Penicillin/Streptomycin, 2% (v/v) B27 supplement with vitamin A (Thermo Fisher Scientific), 1% (v/v) GlutaMAX, 0.5% (v/v) N2 supplement, 1*/*4000 (v/v) insulin (I9278, Sigma-Aldrich), 0.5% (v/v) MEM-NEAA, 1 mg/mL sodium bicarbonate, and 0.07 mg/mL vitamin C.

Organoids were fixed at days 5, 10, 20, 30 and 40 in 4% (w/v) paraformaldehyde (PFA) overnight at 4^°^C, then rinsed three times with PBS (Thermo Fisher Scientific). Fixed organoids were then immobilised in 1% (w/v) agarose for Raman imaging.

To prepare the organoid slices for Raman imaging or immunostaining, the organoid tissues were cryosectioned. Briefly, fixed organoid tissues were first incubated in 30% sucrose at 4^°^C for a day on a roller mixer, then transferred into Tissue-Tek O.C.T. Compound (Sakura Finetek USA), gently swirled, and incubated for 5 minutes. The tissues were then frozen in a cryomold filled with Tissue-Tek O.C.T. For the two cryosections prepared for RS imaging, this was achieved using the technique for high-throughput cryoprocessing previously reported by Reumann et al. ^77^. Compound to form cryoblocks containing the frozen organoids. The cryoblocks were subsequently sectioned into 20 µm slices using a CryoStar NX70 Thermo Cryostat, and collected on positively charged cryoslides. The tissue sections were dried for at least 3 hours before being stored at -20^°^C or used for downstream analysis (fluorescence microscopy and Raman microspectroscopy).

### Immunofluorescence staining and microscopy

For immunostaining, microscopy slices were washed in PBS for 5 minutes to wash off residual OCT, then permeabilised and blocked in PBS containing 0.3% (v/v) Triton X-100 (Thermo Fisher Scientific) and 5% (w/v) BSA (Thermo Fisher Scientific) for 0.5–1.5 hours at room temperature. Primary and secondary antibodies were diluted in PBS containing 0.1% (v/v) Triton X- 100 and 5% (w/v) BSA. Primary antibody solutions were incubated with the slices overnight at 4^°^C, followed by three rinses in PBS, and three washes with PBS-T (PBS+0.01% TX100) for 10–15 minutes on a slow rotating orbital shaker at room temperature. Secondary antibody solutions were then added and incubated for 2 hours at room temperature. Extended information about the primary and secondary antibodies used for immunostaining is provided in Table S1 and Table S2, respectively. Cell membranes were labelled using DiO (BT60011, Biotium) – a fluorescent lipophilic dye – which was added to the secondary antibody solution. DAPI (4’,6-diamidino-2phenylindole) (Thermo Fisher Scientific), diluted 1: 1000 in PBS, was used to stain nuclei for 7 minutes. After this, slides were rinsed three times in PBS, and washed twice in PBS-T and once in PBS for 10–15 minutes on a slowly rotating orbital shaker. Stained organoid slices were then mounted using fluorescent mounting media (S302380-2, Agilent DAKO) or FluoroSave (Millipore). Fluorescent images were acquired using a Leica SP8 confocal microscope and a Nikon TI-2 widefield fluorescent microscope, and processed in FIJI.

### Raman microspectroscopy

Raman spectroscopy measurements were performed using a confocal Raman microscope (alpha300 R, WITec GmbH) equipped with a 532 nm solid-state laser (WITec GmbH). Laser power at the sample plane was set to about 48 mW for organoid cryosections or 32 mW for intact samples. Backscattered light was collected through a water immersion objective – 20× (Zeiss W N-Achroplan, N.A.= 0.5) for organoid cryosections or 63× (Zeiss W Plan-Apochromat, N.A.= 1) for intact samples, and delivered via a 10 µm single-mode silica fiber to an imaging spectrograph (Newton, Andor Technology Ltd) with a 600 groove/mm grating and a thermoelectrically cooled CCD detector (− 60^°^C). The system provided a spectral resolution of ∼2–3 cm^−1^ over the wavenumber range 0–3670 cm^−1^. Spectra were acquired with pixel sizes in *x* and *y* and step sizes in *z* ranging from 1 µm to 10 µm, and an integration time of 0.5 seconds.

### Spectral preprocessing of Raman imaging data

Acquired Raman spectroscopy measurements were exported as MATLAB files using the WITec Project FIVE Suite software and loaded in Python using RamanSPy ^54^. Following this, each scan was individually preprocessed using RamanSPy as follows: (1) spectral cropping whereby wavenumbers less than 300 cm^*−*1^ were removed; (2) cosmic rays removal, using the algorithm reported by Whitaker and Hayes ^78^, with a kernel size *m* = 3 and *z*-value threshold *τ* = 6; (3) denoising with a third-order Whittaker smoother ^79^ with *λ* = 10^3^; (4) baseline correction with asymmetric least squares ^80^ with *λ* = 10^5^; and (5) vector normalisation. Normalisation was applied on a pixel-wise basis for our timepoint data to account for depth-dependent signal attenuation. For the rest of the data, global vector normalisation was performed where intensity values were divided by the *ℓ*^2^-norm of the spectrum with the highest norm in the given scan. Additionally, wavenumber calibration was performed on our cryosection scans by calculating the median position of the laser peak (Rayleigh scattering) and setting it to zero.

### Hyperspectral unmixing analysis

Chemometric analysis was performed using a custom autoencoder neural network model designed for blind hyperspectral unmixing. In this context, Raman measurements **x**_*i*_ ∈ ℝ^*b*^ are treated as mixtures **x**_*i*_ = ℱ (*M, α*_*i*_) of a set of *n* unknown endmember components *M* ∈ ℝ^*n*×*b*^ based on their relative abundances *α*_**i**_ ∈ ℝ^*n*^ in a given measurement **x**_*i*_. Blind unmixing aims to decompose a set of Raman measurements 𝒳 ∈ ℝ^*m*×*b*^ (i.e. a given scan) into endmember components *M* and their relative abundances *A* ∈ ℝ^*m*×*n*^.

To achieve this, we train an autoencoder model 𝒜 consisting of an encoder module ℰ and a decoder module 𝒟. The encoder was responsible for mapping input spectra **x** into latent space representations **z** = ℰ (**x**), and the decoder was responsible for mapping these latent representations into reconstructions of the original input 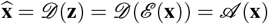. The model was trained in an unsupervised manner by minimising the reconstruction error between the input **x** and the output 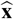. During this process, the model was guided to learn the endmember components *M* and their relative abundances *A* through the introduction of related physical constraints. Below, we provide additional details about the developed architecture and training procedure. For more information about hyperspectral unmixing, the reader is pointed to previous works by Keshava and Mustard ^55^ and Li et al. ^56^. For more information about autoencoders, the reader is pointed to the work of Goodfellow et al. ^81^.

The encoder ℰ comprised two separate blocks applied sequentially. The first part was a multi-branch convolutional block comprising four parallel convolutional layers with kernel sizes of 5, 10, 15 and 20, designed to capture patterns at multiple spectral scales. Each convolutional layer contained 32 filters with ReLU activation, He initialisation ^82^ and ‘same’ padding. Batch normalisation ^83^ and dropout with a rate of 0.2^84^ were applied to each convolutional layer to improve training stability and generalisation. The outputs of the four convolutional layers were merged channel-wise through a fully connected layer to yield an output of dimension matching that of the input spectrum. The rationale behind this was to transform intensity values into representations that capture local spectral features (e.g. peak shape, width, local neighbourhood) and thus promote better generalisability. The second part of the encoder was a fully connected dimensionality reduction block, applied to learn patterns between the learnt spectral features. This block comprised a series of fully connected layers of sizes 256, 128, 64 and 32 with He initialisation and ReLU activation. Batch normalisation and dropout (rate of 0.5) were also applied at each fully connected layer. The block was followed by a final fully connected layer (Xavier uniform initialisation ^85^) that reduced the final 32 features to a latent space of size *n*. The number *n* was treated as a hyperparameter that encodes the number of endmembers to extract, with latent representations treated as abundance fractions. To improve c was enforced in the latent space using a ‘softly-rectified’ hyperbolic tangent function 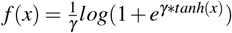, with *γ* = 10, as we previously reported ^53^. This ensured that latent values were constrained to the range (0, 1) in accordance with the abundance non-negativity constraint, but does not enforce abundances to sum to one (abundance sum-to-one constraint).

The decoder 𝒟 consisted of a single fully connected layer (Xavier uniform initialisation) mapping the *n*-dimensional latent space representations back to the original spectral dimension *b*. In this layer, we used a linear activation and no bias term. Under this setup, the decoder mimics linear unmixing, where endmember signatures are encoded in the learnt weight matrix of the decoder layer ^53^. A non-negative kernel constraint was used to enforce the non-negativity of endmember signatures using weight clipping during training. To accelerate model training, the weight matrix in the linear decoder layer was initialised before training using endmember signatures derived with vertex component analysis ^64^.

Autoencoder models were trained for 5 epochs using the Adam optimiser ^86^ with a batch size of 32. The learning rate was set to 0.001 for our two cryosection scans and subcellular day-10 scan, or 0.003 for all other data. An exponential learning rate decay was used to ensure stable convergence, applying a factor of 0.9 to the learning rate after each epoch. The training loss ℒ was defined as a weighted sum of the mean squared error (MSE) and spectral angle divergence (SAD) between input spectra **x** and output reconstructions 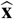:

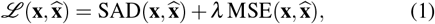

where

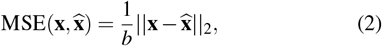

with *b* denoting the dimension of **x** (i.e. the number of spectral bands), and

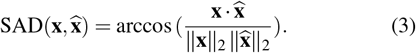

The weighting factor *λ* was set to 1000.

The number of endmembers to derive was estimated individually for each scan/dataset of interest using principal component analysis. Specifically, we computed the cumulative explained variance as a function of the number of principal components and manually selected the number of components at the point where the increase in explained variance visually plateaued.

Autoencoder model implementation, training and evaluation steps were carried out using TensorFlow ^87^. Analysis with baseline chemometric methods was performed using RamanSPy ^54^.

### Timepoint analysis

Quantitative analysis is based on Raman imaging data from three batches of cerebral organoids, cultured in independent experiments using stem cells at different passage numbers. For our quantification analysis, manual background segmentation was performed using the derived abundance reconstructions as a reference. The obtained binary masks were used to (i) discard background pixels by setting abundance values in the background area to zero for all channels, and (ii) estimate the relative tissue portion in each scan. Scans with an organoid area below 30% were not used for the subsequent quantification to avoid potential biasing towards components in the outer tissue layers. In total, six scans out of the available 72 were discarded (Figure S14B). Finally, the mean abundance of endmembers was calculated for each scan and normalised by the organoid area. Representative abundance images were produced without background removal, after scaling the data by a factor of 2 in *z* (organoid depth) to achieve an equal effective pixel size of 5 µm in *x*-*y* and *z*, and thus provide a more accurate depiction of the organoid morphology and dimensions.

## Data and Code Availability

The research data and code supporting this paper will be made available upon publication.

## Acknowledgments

The authors thank Dr Simon Vilms Pedersen for discussions, Kevin Giraldo Rodriguez for assistance with Raman imaging of an organoid cryosection, Ellena O’Keeffe for performing immunostaining of an organoid cryosection, and Dr Akemi Nogiwa Valdez for data management support. The authors also thank the Oxford Organoid Hub (OOH) for providing imaging infrastructure. Cryosectioned organoids imaged by fluorescence microscopy and Raman microspectroscopy for correlative analysis were generated by D.R. in the Knoblich laboratory (Vienna, Austria). D.G. acknowledges support by UK Research and Innovation (UKRI Centre for Doctoral Training in AI for Healthcare, grant number EP/S023283/1). R.X. and M.M.S. acknowledge support from the Engineering and Physical Sciences Research Council (EP/P00114/1 and EP/T020792/1). D.R. and M.M.S. acknowledge support from UK Research and Innovation (UKRI) Innovate UK under the UK government’s Horizon Europe guarantee (10145749) associated to a grant from the European Union’s Horizon Europe research and innovation programme (101071203). M.M.S. and D.R. acknowledge support from the Advanced Research and Invention Agency (SCNI-PR01-P11). Á.F.-G. acknowledges support from the Schmidt Science Fellows, in partnership with the Rhodes Trust.

M.B. acknowledges support from the Engineering and Physical Sciences Research Council (EP/N014529/1, funding the EPSRC Centre for Mathematics of Precision Healthcare at Imperial College London, and EP/T027258/1). M.M.S. acknowledges support from the Royal Academy of Engineering Chair in Emerging Technologies award (CiET2021\94). Figures were assembled in BioRender.

## Author Contributions

D.G. and R.X. contributed equally to this work. D.G., R.X., D.R., Á.F.-G., M.B. and M.M.S. contributed to research design and conceptualisation. D.G. developed the computational pipeline, implemented software and performed data analysis. R.X., D.R. and X.Z. contributed to organoid culture and Raman imaging. R.X. and D.R. contributed to sample preparation and fluorescence microscopy. D.G., R.X., D.R., Á.F.-G., M.B. and M.M.S. contributed to data interpretation. M.B. and M.M.S. supervised the study. D.G. wrote the paper, with input from all authors. All authors have approved the final version of the manuscript.

## Supplementary Information

### Supplementary Information includes

Supplementary Tables S1–S2.

Supplementary Figures S1–S15.

**Figure S1:**
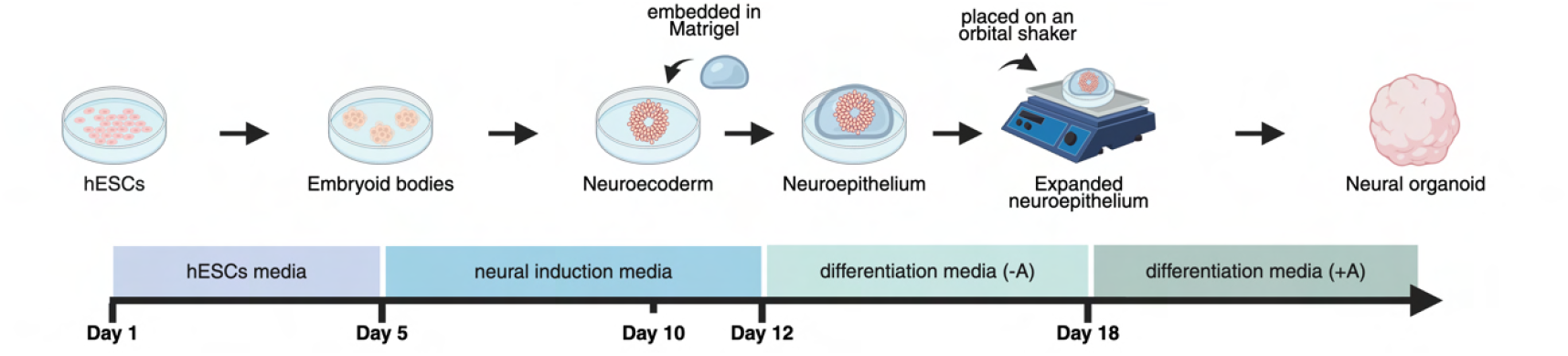
Schematic diagram of organoid culture. Neural organoids were grown using a protocol for unguided differentiation based on previous work from Lancaster et al. ^7^.

**Table S1:** Primary antibodies. Extended information about the primary antibodies used for immunofluorescence staining. ‘Cat. no.’ stands for ‘Catalogue number’.

| Figure panel(s) | Antigen | Species | Producer | Cat. no. | Dilution |
| --- | --- | --- | --- | --- | --- |
| Figure 1C | SOX2 | Rabbit | Abcam | ab97959 | 1:600 |
| | PKCζ | Mouse | Santa Cruz Biotechnology | sc-17781 | 1:200 |
| Figure 2C, Figure S3B | SOX2 | Goat | R&D Systems | AF2018 | 1:100 |
|  | TUJ1 | Mouse | BioLegend | 801201 | 1:100 |
| Figure 2D, Figure S5B | SOX2 | Goat | R&D Systems | AF2018 | 1:100 |
|  | TUJ1 | Mouse | BioLegend | 801201 | 1:100 |
| Figure 4A | SOX2 | Rabbit | Abcam | ab97959 | 1:600 |
| | PKCζ | Mouse | Santa Cruz Biotechnology | sc-17781 | 1:200 |
|  | MAP2 | Chicken | Abcam | ab5392 | 1:500 |

**Table S2:**
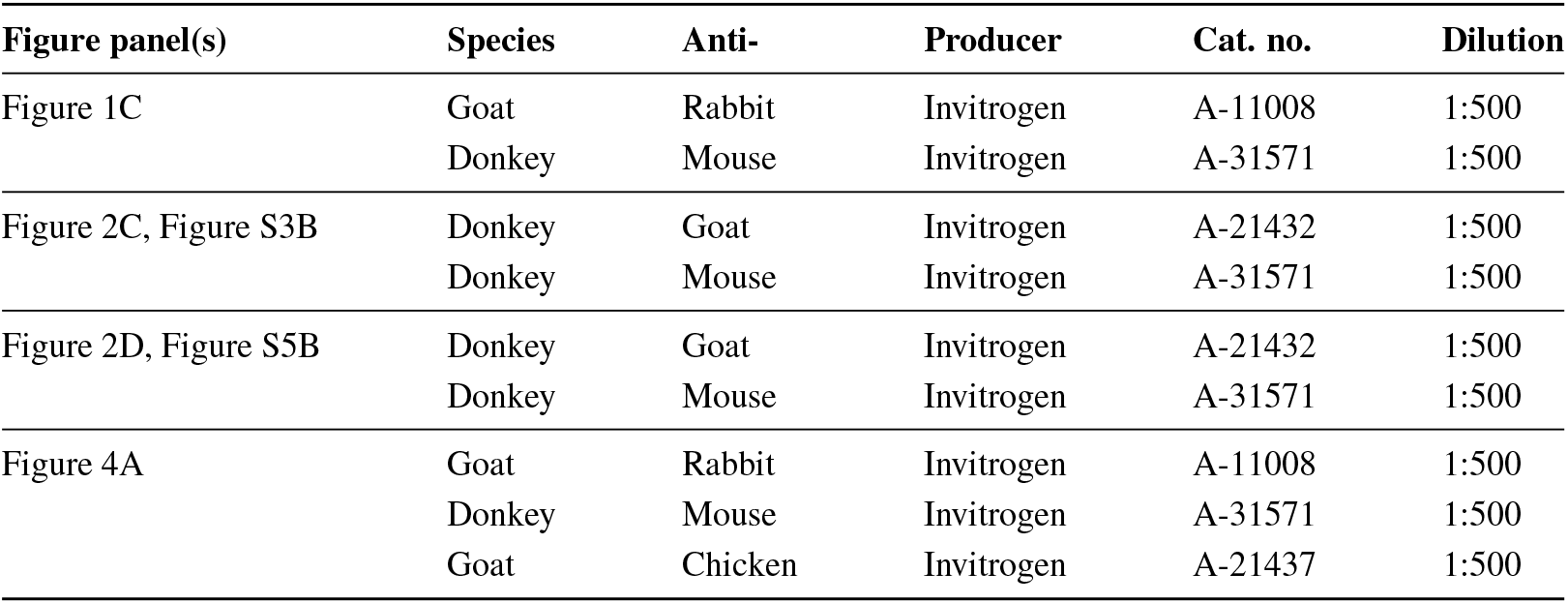
Secondary antibodies. Extended information about the secondary antibodies used for immunofluorescence staining. ‘Cat. no.’ stands for ‘Catalogue number’.

| Figure panel(s) | Species | Anti- | Producer | Cat. no. | Dilution |
| --- | --- | --- | --- | --- | --- |
| Figure 1C | Goat | Rabbit | Invitrogen | A-11008 | 1:500 |
|  | Donkey | Mouse | Invitrogen | A-31571 | 1:500 |
| Figure 2C, Figure S3B | Donkey | Goat | Invitrogen | A-21432 | 1:500 |
|  | Donkey | Mouse | Invitrogen | A-31571 | 1:500 |
| Figure 2D, Figure S5B | Donkey | Goat | Invitrogen | A-21432 | 1:500 |
|  | Donkey | Mouse | Invitrogen | A-31571 | 1:500 |
| Figure 4A | Goat | Rabbit | Invitrogen | A-11008 | 1:500 |
|  | Donkey | Mouse | Invitrogen | A-31571 | 1:500 |
|  | Goat | Chicken | Invitrogen | A-21437 | 1:500 |

**Figure S2:**
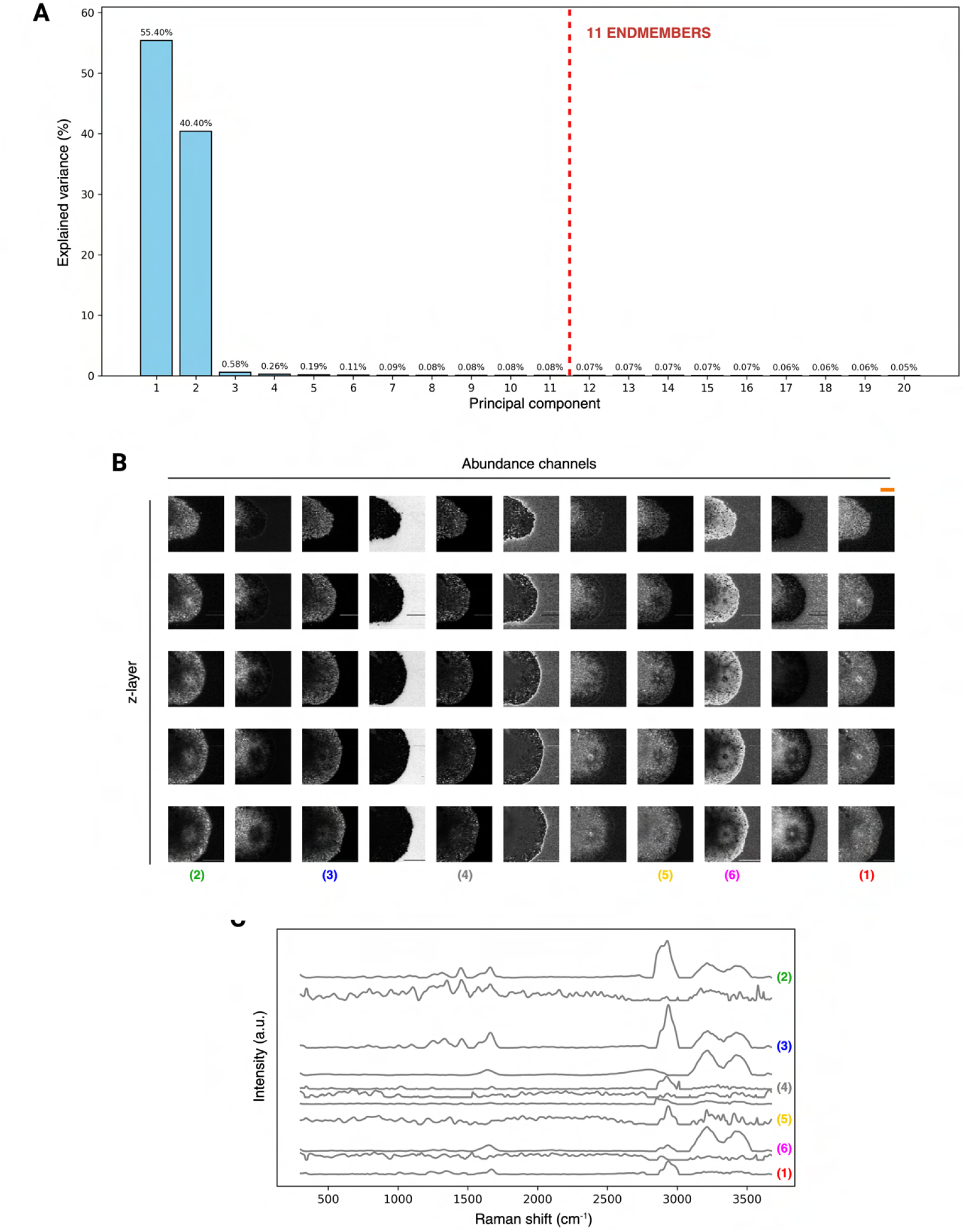
Extended unmixing results for low-resolution volumetric Raman imaging scan of an intact day-20 neural organoid (as shown in Figures 1D–H in the main text). **A**, Principal component analysis of preprocessed RS data. The number of endmember components to extract was set to 11. **B**, Derived abundance images (normalised for visualisation purposes). Scale bar: 100 µm (top right, marked in orange). **C**, Corresponding endmembers. Endmember and abundance numbering indicate the components presented in the main text.

**Figure S3:**
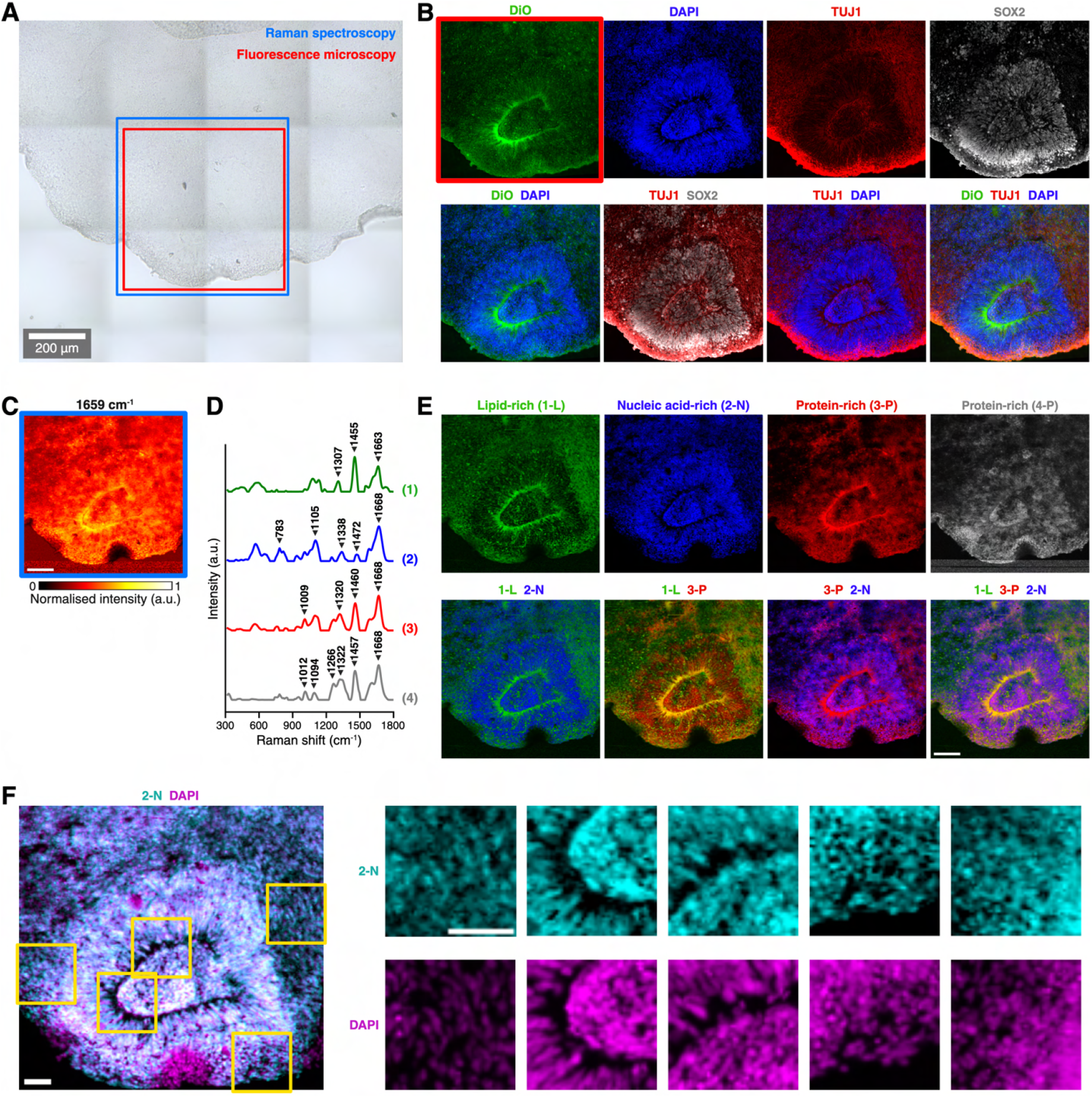
Extended results for Raman imaging scan of a first day-30 neural organoid cryosection (as shown in Figures 2B and 2C in the main text). **A**, Brightfield image of a sectioned neural organoid tissue region imaged using Raman spectroscopy (blue) and fluorescence microscopy (red). Scale bar: 200 µm. **B**, Fluorescence microscopy images with markers for nuclei (DAPI), membranes (DiO), neural progenitors (SOX2) and neurons (TUJ1). **C**, Raman intensity profile at the 1659 cm^−1^ band (linked to proteins and lipids). Image size: 300 × 300 pixels. Step size: 2 µm. Scale bar: 100 µm. **D–E**, Result of hyperspectral unmixing analysis. **D**, Derived tissue-specific endmember signatures. Endmembers scaled for visualisation purposes (maximum intensity set to 1). **E**, Corresponding abundance maps. Scale bar: 100 µm. Endmember 1 (green) presented a lipid-rich spectral signature with distinct peaks at 1307 cm^−1^, 1455 cm^−1^, and 1663 cm^−1^. Its abundance was consistent with membrane structures, including cellular membranes and the apical domain of the visible neural rosette. The signal of this endmember showed a high degree of visual overlap with DiO and TOJ1. Endmember 2 (blue) displayed a nucleic acid-rich profile with a prominent band at 783 cm^−1^, alongside vibrations linked to proteins (1338 cm^−1^, 1472 cm^−1^ and 1668 cm^−1^). Strong glass signal was also observed around 570 cm^−1^ and 1100 cm^−1^. The abundance map revealed well-defined, spherical compartments, closely resembling cell nuclei, with increased density within the present neural rosette structure. Endmembers 3 (red) and 4 (grey) shared similar spectral signatures, dominated by protein-associated peaks around ∼1010 cm^−1^, ∼1100 cm^−1^, ∼1265 cm^−1^, ∼1320 cm^−1^, ∼1460 cm^−1^, and ∼1670 cm^−1^. Both endmembers were generally broadly distributed across the sample. Endmembers 3 showed increased signal at the apical domain of the present rosette, potentially suggesting structural proteins. The abundance of endmember 4 was increased within the rosette structure. **F**, Overlaid images of DAPI and endmember associated with nucleic acids (2-N). Scale bars: 25 µm.

**Figure S4:**
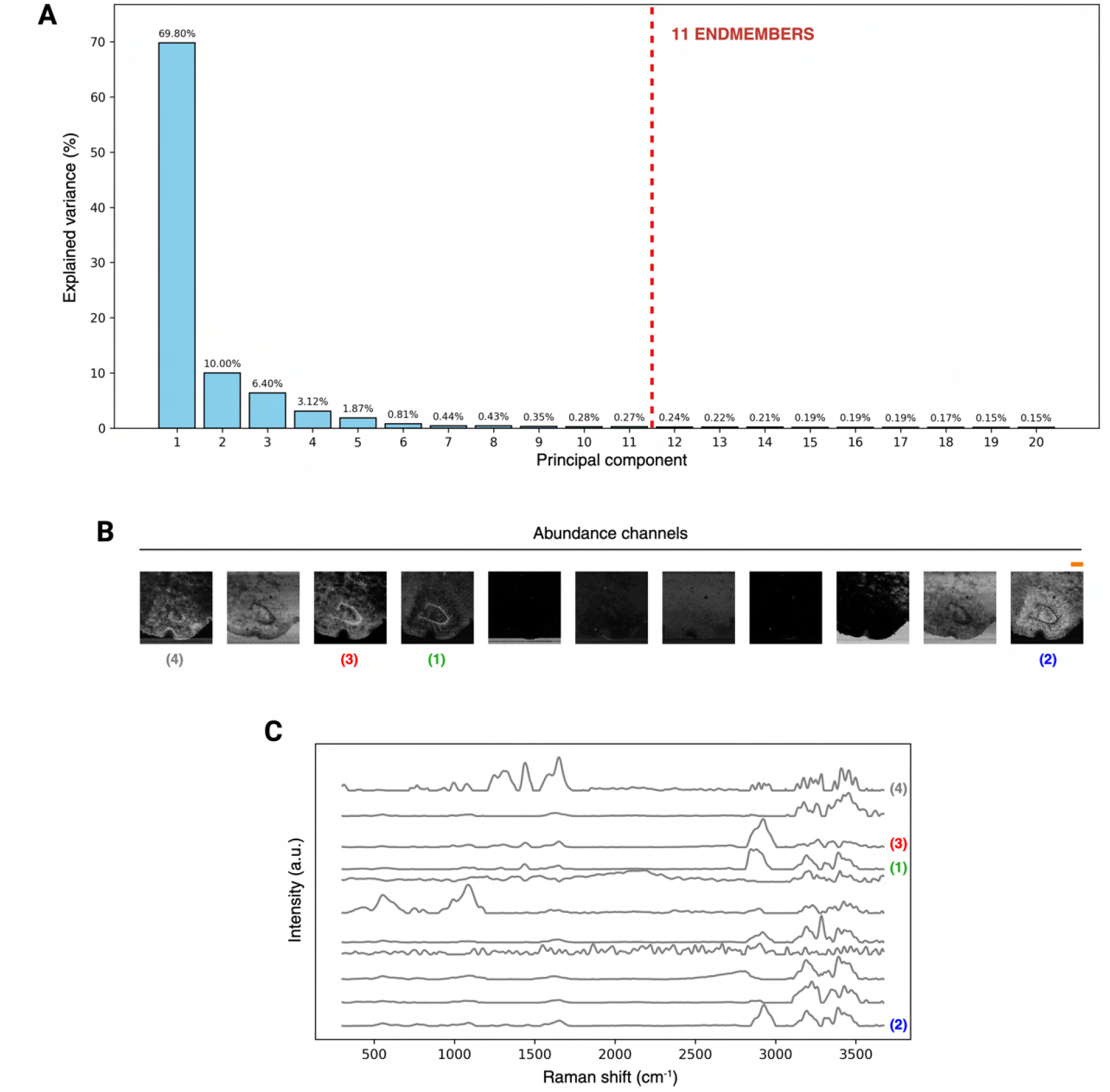
Extended unmixing results for Raman imaging scan of a day-30 neural organoid cryosection (as shown in Figures 2B and 2C in the main text). **A**, Principal component analysis of preprocessed RS data. The number of endmember components to extract was set to 11. **B**, Derived abundance images (normalised for visualisation purposes). Scale bar: 100 µm (top right, marked in orange). **C**, Corresponding endmembers. Endmember and abundance numbering indicate the components presented in the main text.

**Figure S5:**
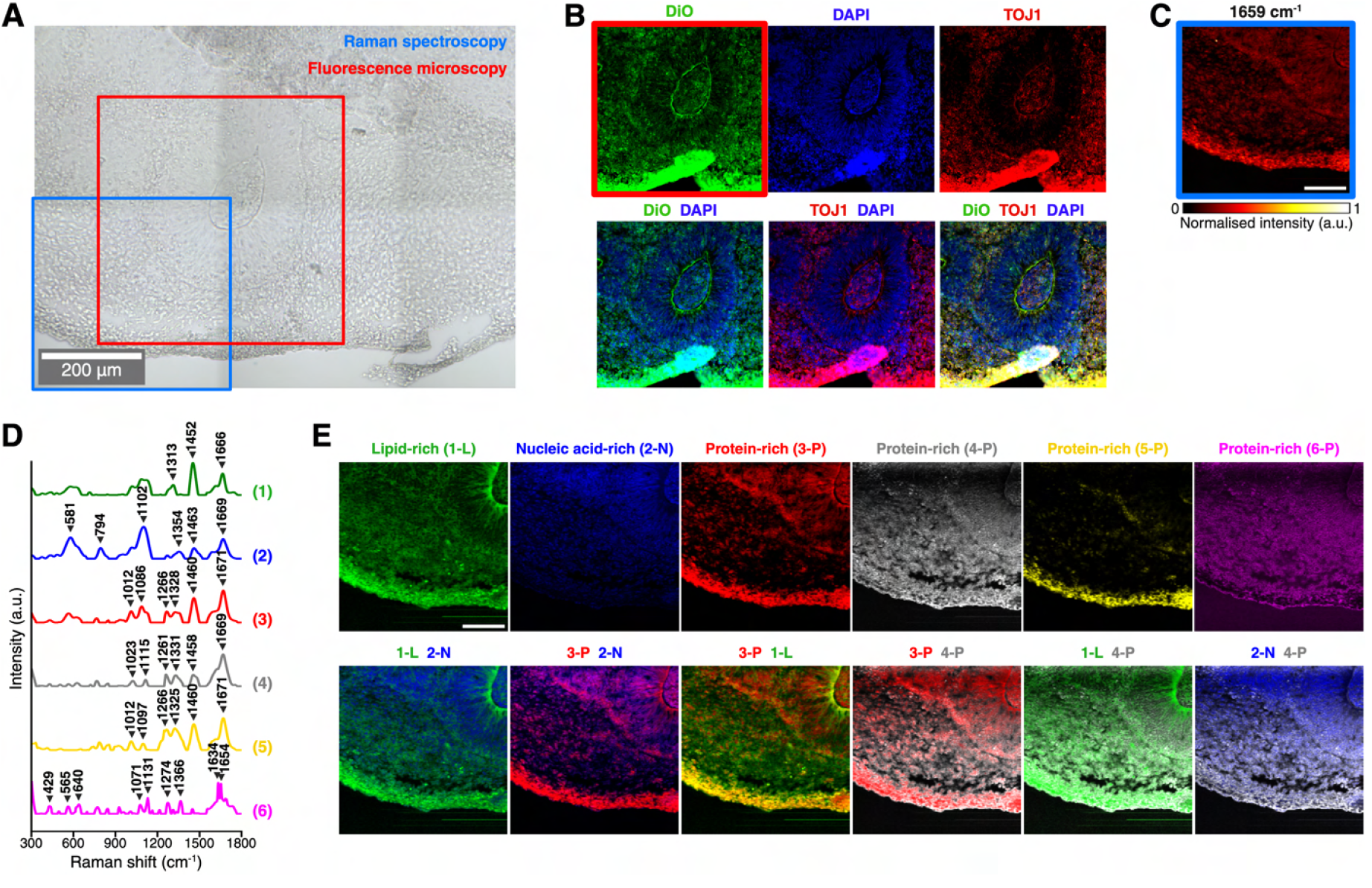
Extended results for Raman imaging scan of a second day-30 neural organoid cryosection (as shown in Figures 2B and 2D in the main text). **A**, Brightfield image of a sectioned neural organoid tissue imaged using Raman spectroscopy (blue) and fluorescence microscopy (red). Scale bar: 200 µm. **B**, Fluorescence microscopy images with markers for nuclei (DAPI), membranes (DiO) and neurons (TUJ1). Due to partial tissue detachment during the staining process, a localised increase in fluorescence intensity was observed at the tissue edge. This peripheral region was identified as an artefact and was therefore excluded from subsequent analyses. **C**, Raman intensity profile at the 1659 cm^−1^ band (linked to proteins and lipids). Image size: 200 × 200 pixels. Step size: 2 µm. Scale bar: 100 µm. **D–E**, Result of hyperspectral unmixing analysis. **D**, Derived tissue-specific endmember signatures. Endmembers scaled for visualisation purposes (maximum intensity set to 1). **E**, Corresponding abundance maps. Scale bar: 100 µm. Endmember 1 (green) exhibited a lipid-rich profile with prominent peaks at 1313 cm^−1^, 1452 cm^−1^, and 1666 cm^−1^. The corresponding abundance map presented morphological architecture consistent with membrane-like structures, including individual cellular membranes and the apical domain of the present neural rosette. Endmember 2 (blue) displayed a strong nucleic acid band at 794 cm^−1^, alongside several vibrations linked to protein species (1354 cm^−1^, 1463 cm^−1^ and 1669 cm^−1^), as well as bands attributed to the glass slide at (581 cm^−1^ and 1102 cm^−1^). The abundance map revealed well-defined, spherical compartments, closely resembling cell nuclei. Endmembers 3 (red), 4 (grey) and 5 (yellow) exhibited similar spectral signatures, dominated by protein-associated peaks around ∼1015 cm^−1^, ∼1100 cm^−1^, ∼1265 cm^−1^, ∼1330 cm^−1^, ∼1460 cm^−1^, and ∼1670 cm^−1^. Endmember 4 showed a broad spatial distribution, likely corresponding to cytoplasmic or general background tissue signal. Endmembers 3 and 5, which showed a more pronounced 1460 cm^−1^ band, and an increased signal near the edge of the tissue section, likely due to tissue curling, as well as the basal domains of the present neural rosettes. Endmember 6 (magenta) presented a noisier signature, with prominent protein-associated peaks at 1131 cm^−1^ (C–N stretching), 1274 cm^−1^, and 1366 cm^−1^ (tryptophan). Its spatial distribution showed substantial overlap with that of Endmember 4.

**Figure S6:**
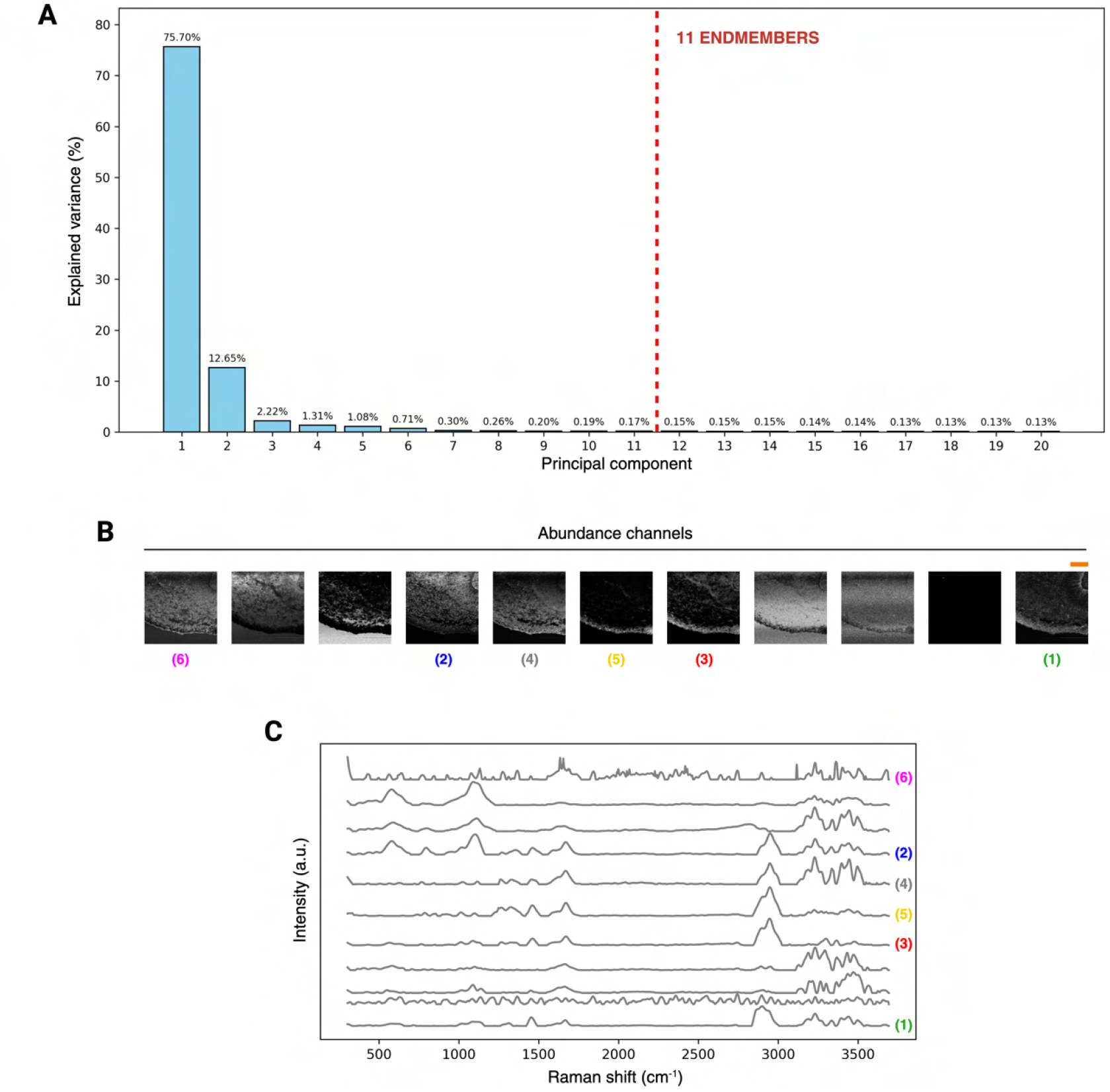
Extended unmixing results for Raman imaging scan of a second day-30 neural organoid cryosection (as shown in Figures 2B and 2D in the main text). **A**, Principal component analysis of preprocessed RS data. The number of endmember components to extract was set to 11. **B**, Derived abundance images (normalised for visualisation purposes). Scale bar: 100 µm (top right, marked in orange). **C**, Corresponding endmembers. Endmember and abundance numbering indicate the components presented in the main text.

**Figure S7:**
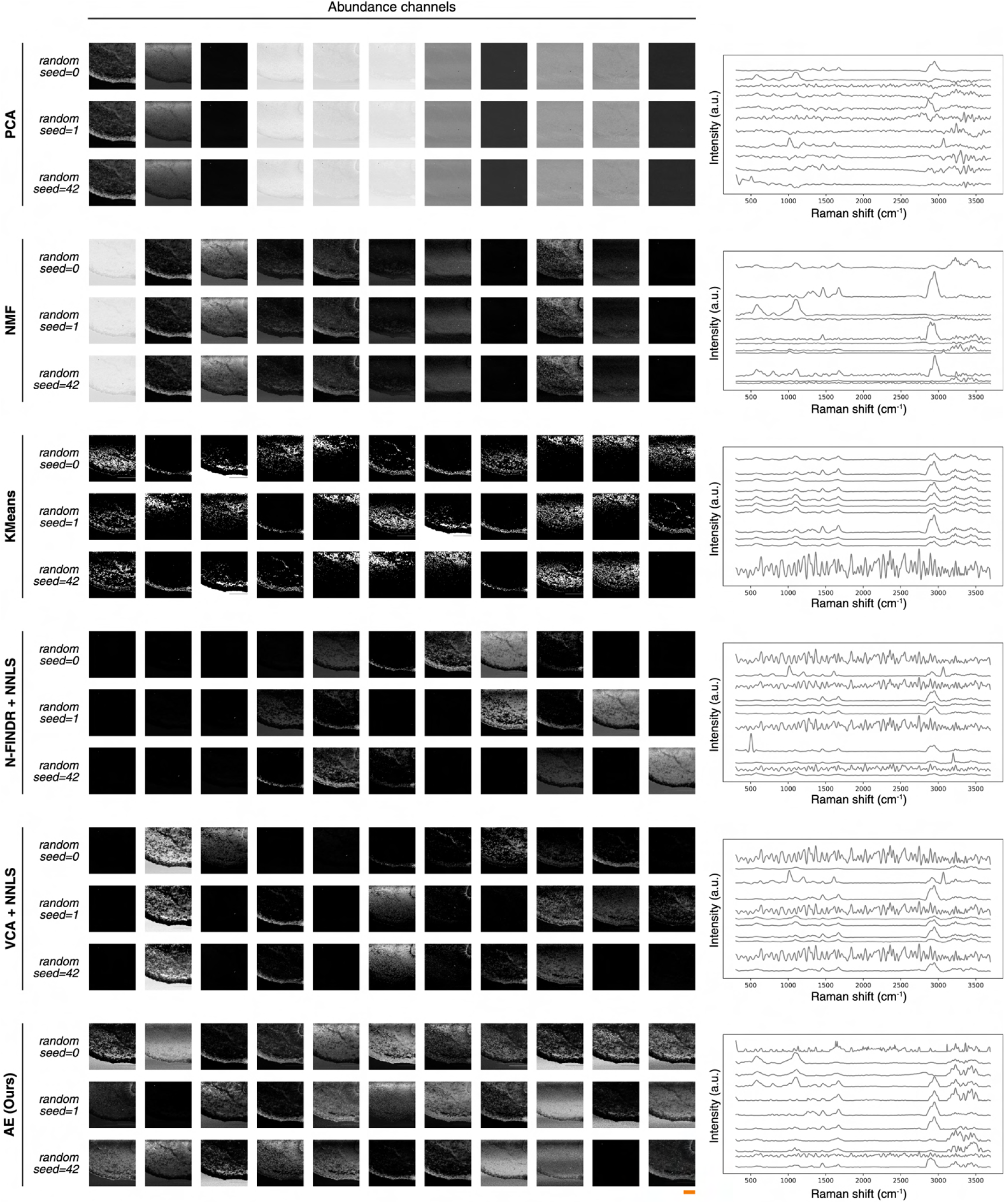
Comparative analysis of different chemometric methods on Raman imaging scan of a day-30 neural organoid cryosection (as shown in Figures 2B and 2D in the main text). Methods include: principal component analysis (PCA), non-negative matrix factorisation (NMF), k-means clustering, N-FINDR + non-negative least squares (NNLS), vertex component analysis (VCA) + NNLS, our autoencoder (AE) pipeline. Abundance images were normalised for visualisation purposes. Endmember signatures displayed on the right correspond to the analyses performed with a random seed set to 42. PCA yielded a single tissue-specific component – its first principal component – that effectively captured the overall tissue architecture. The remaining components displayed convoluted abundances with high uniform background signals and negligible biochemical specificity. In contrast, NMF identified three distinct endmembers, resembling major molecular species such as protein-rich, lipid-rich and nucleic acid-rich components. However, it was unable to resolve finer biochemical features beyond these primary categories. K-means clustering showed limited biochemical interpretability, segmenting the data into different tissue regions, possibly due to intensity variation instead of meaningful compositional components. More specialised unmixing techniques like N-FINDR+NNLS exhibited serious susceptibility to artefacts, producing multiple abundance maps with isolated “noisy” pixels. Beyond one general tissue-level component, N-FINDR+NNLS failed to resolve distinct cellular structures such as nuclei or membranes. VCA+NNLS demonstrated improved performance relative to N-FINDR+NNLS, producing five distinct abundance maps that captured features such as nuclei, the apical domain of the imaged neural rosette and potentially cytoplasmic regions. Nonetheless, some artefacts and shading inconsistencies persisted – for instance, within the nuclear component (channel 6), which displayed a strong signal in the upper field that markedly diminished toward the bottom. Scale bar: 100 µm (bottom right, marked in orange).

**Figure S8:**
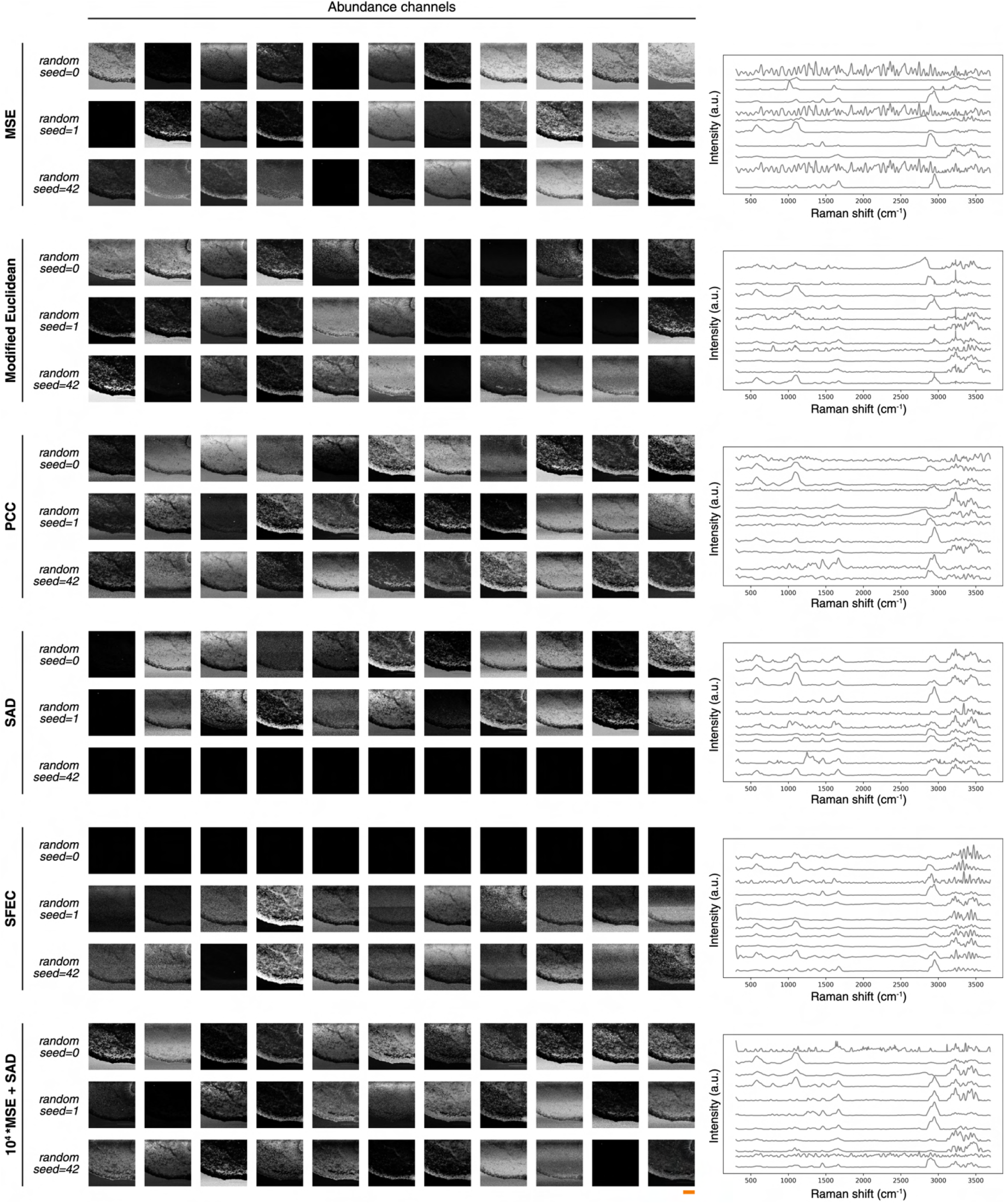
Effect of training loss on autoencoder performance on Raman imaging scan of a day-30 neural organoid cryosection (as shown in Figures 2B and 2D in the main text). Losses tested include: mean squared error (MSE), modified Euclidean ^88^, Pearson’s correlation coefficient (PCC), spectral angle divergence (SAD) ^57^, squared first-difference Euclidean cosine (SFEC) ^89^, and our custom loss based on a combination of MSE and SAD. Abundance images were normalised for visualisation purposes. Endmember signatures displayed on the right correspond to the analyses performed with a random seed set to 42. Scale bar: 100 µm (bottom right, marked in

**Figure S9:**
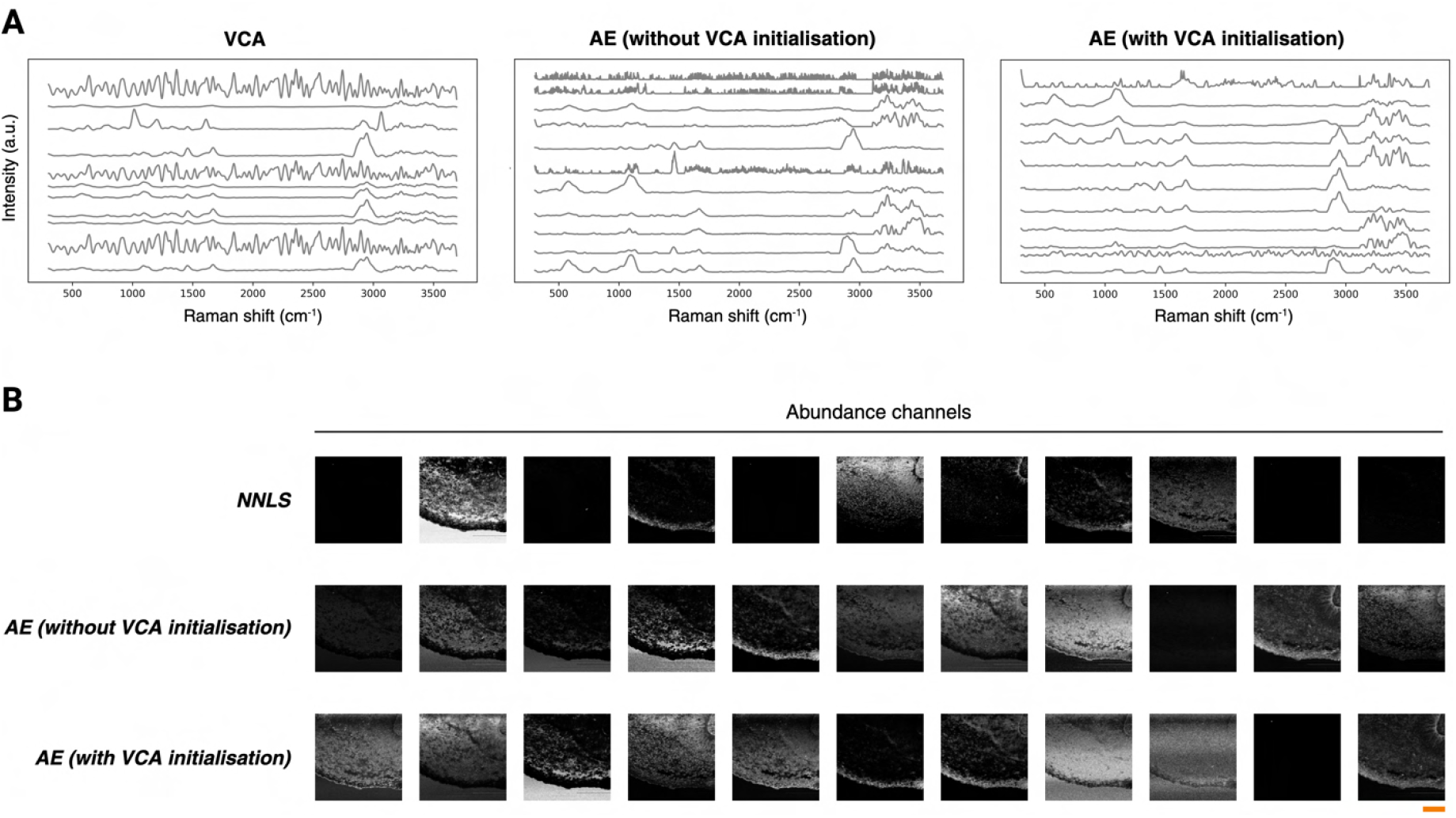
Effect of endmember initialisation on autoencoder performance on Raman imaging scan of a day-30 neural organoid cryosection (as shown in Figures 2B and 2D in the main text). **A**, Derived endmembers with (i) vertex component analysis (VCA), (ii) our autoencoder model without VCA initialisation, and (iii) our autoencoder model with VCA initialisation. **B**, Derived abundances with (i) non-negative least squares (NNLS) performed using the VCA endmembers displayed in **A**, (ii) our autoencoder model without VCA initialisation, and (iii) our autoencoder model with VCA initialisation. Abundance images were normalised for visualisation purposes. Scale bar: 100 µm (bottom right, marked in orange).

**Figure S10:**
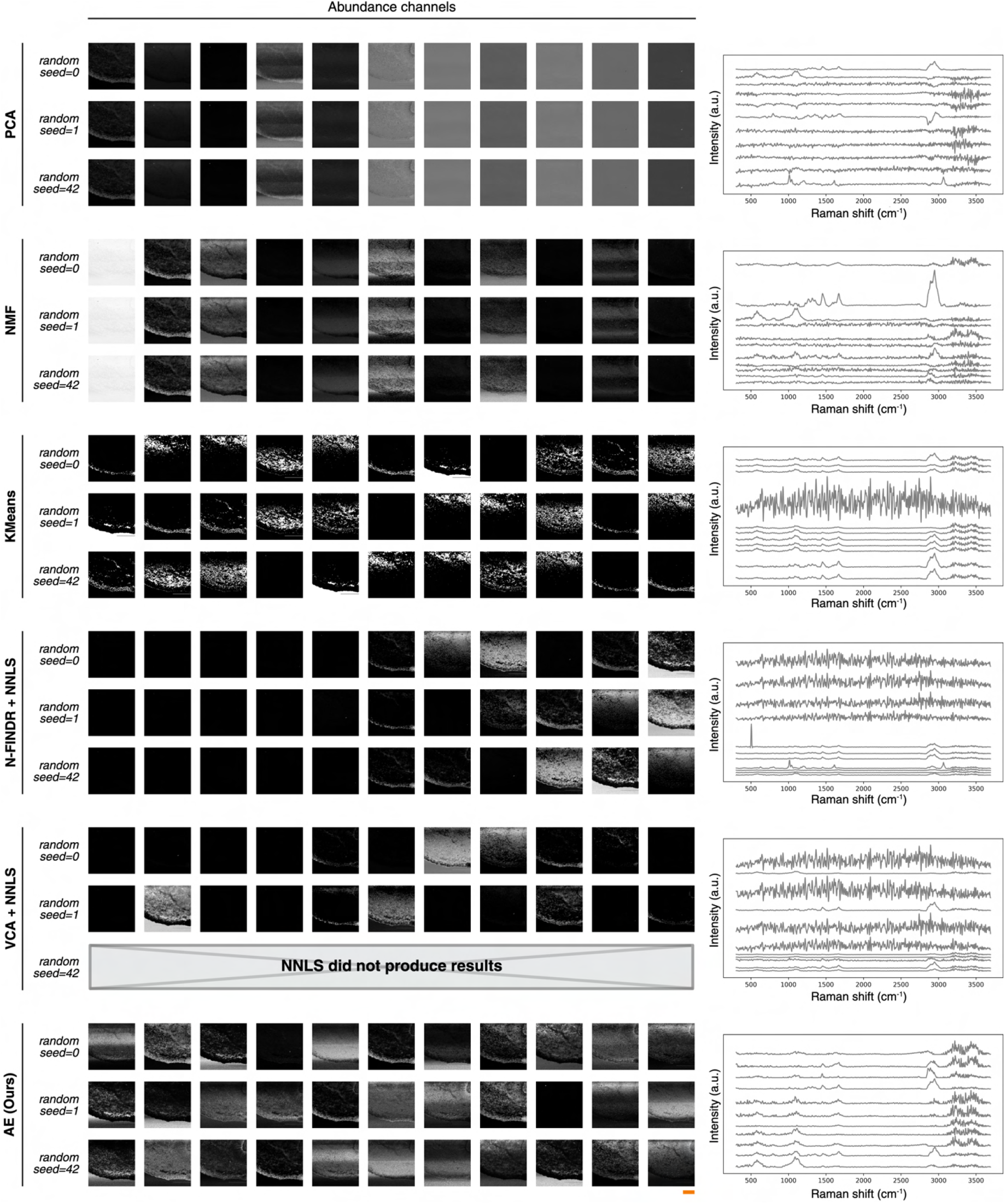
Comparative analysis of different chemometric methods on Raman imaging scan of a day-30 neural organoid cryosection in the presence of higher degrees of experimental noise (as shown in Figures 2B and 2D in the main text). Higher noise levels were simulated by applying a more conservative denoising procedure with a second-order Savitzky-Golay filter with a window size of 7^90^. Methods include: principal component analysis (PCA), non-negative matrix factorisation (NMF), k-means clustering, N-FINDR + non-negative least squares (NNLS), vertex component analysis (VCA) + NNLS, our autoencoder (AE) pipeline. One of the analyses with VCA+NNLS failed to converge, with the NNLS implementation terminating after exceeding the maximum number of iterations. Abundance images were normalised for visualisation purposes. Endmember signatures displayed on the right correspond to the analyses performed with a random seed set to 42. Scale bar: 100 µm (bottom right, marked in orange).

**Figure S11:**
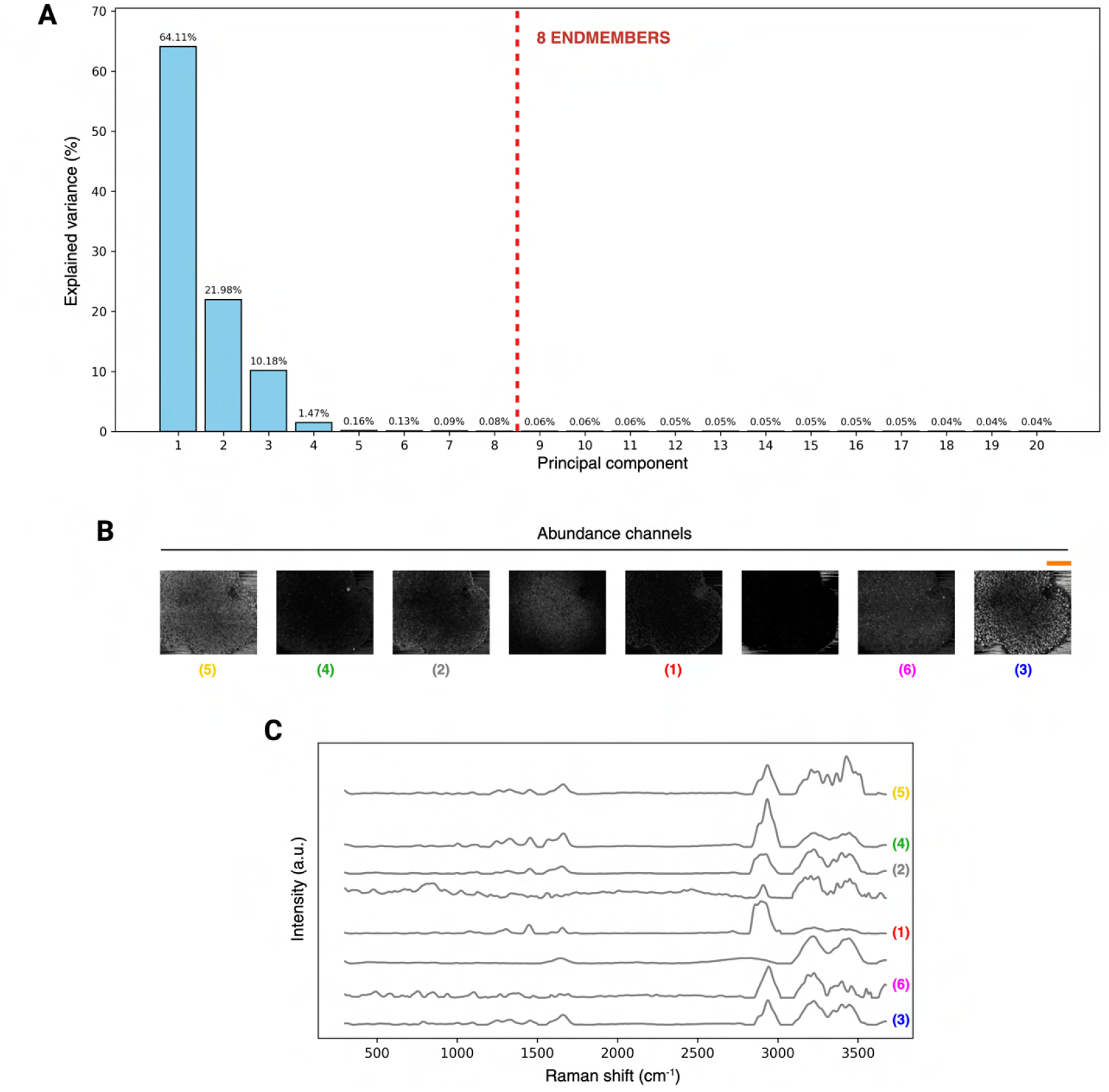
Extended unmixing results for high-resolution Raman imaging scan of an intact day-10 neural organoid (as shown in Figures 3B–E in the main text). **A**, Principal component analysis of preprocessed RS data. The number of endmember components to extract was set to 8. **B**, Derived abundance images (normalised for visualisation purposes). Scale bar: 100 µm (top right, marked in orange). **C**, Corresponding endmembers. Endmember and abundance numbering indicate the components presented in the main text.

**Figure S12:**
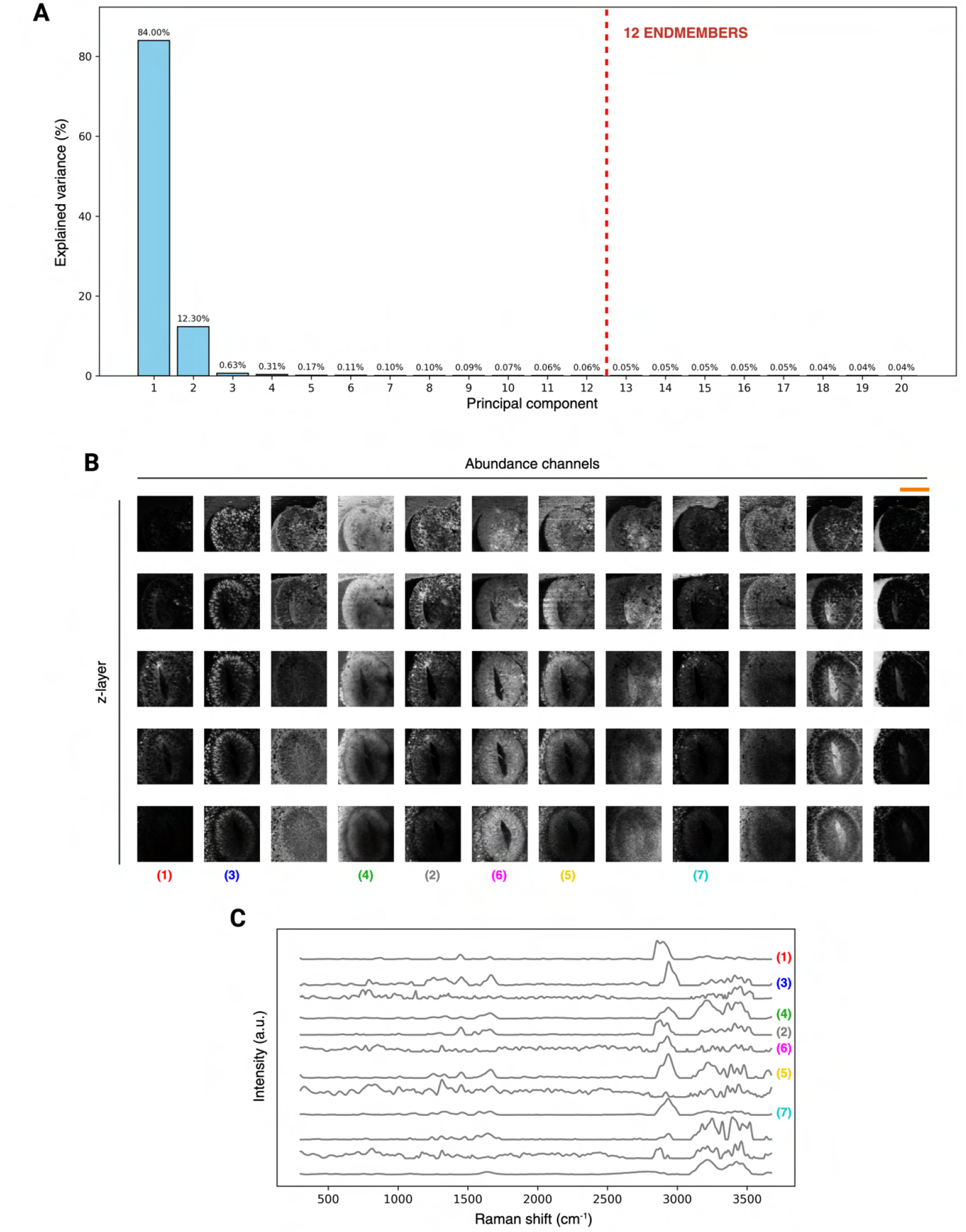
Extended unmixing results for volumetric Raman imaging scan of a neural rosette within an intact day-20 neural organoid (as shown in Figures 4B–E in the main text). **A**, Principal component analysis of preprocessed RS data. The number of endmember components to extract was set to 12. **B**, Derived abundance images (normalised for visualisation purposes). Scale bar: 100 µm (top right, marked in orange). **C**, Corresponding endmembers. Endmember and abundance numbering indicate the components presented in the main text.

**Figure S13:**
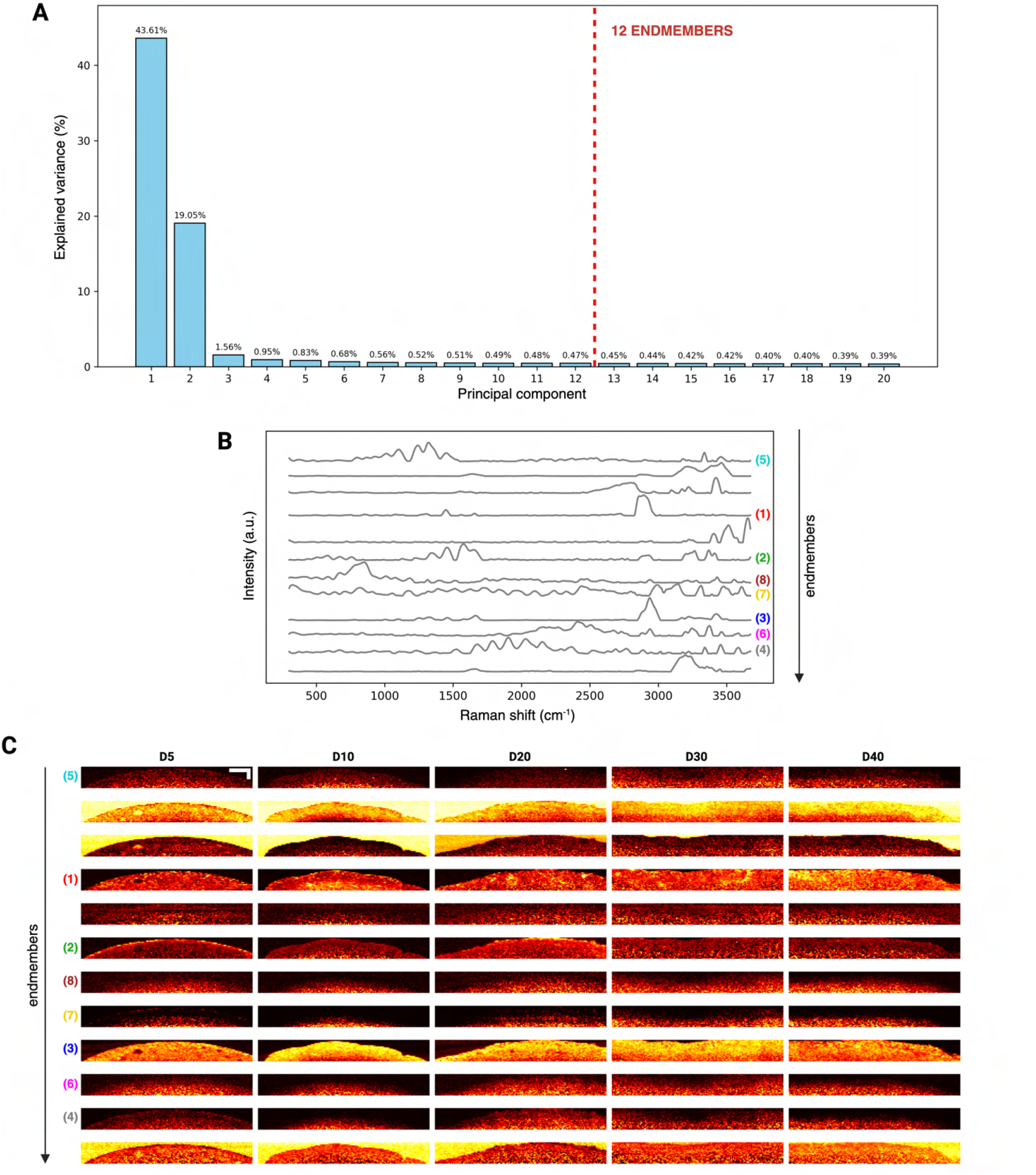
Extended unmixing results for timepoint Raman imaging data (as shown in Figures 5B–D in the main text). **A**, Principal component analysis of preprocessed RS data. The number of endmember components to extract was set to 12. **B**, Derived endmembers. **C**, Corresponding abundance images (normalised for visualisation purposes) for five representative organoid samples. Endmember and abundance numbering indicate the components presented in the main text. Scale bar: 100 µm (top left).

**Figure S14:**
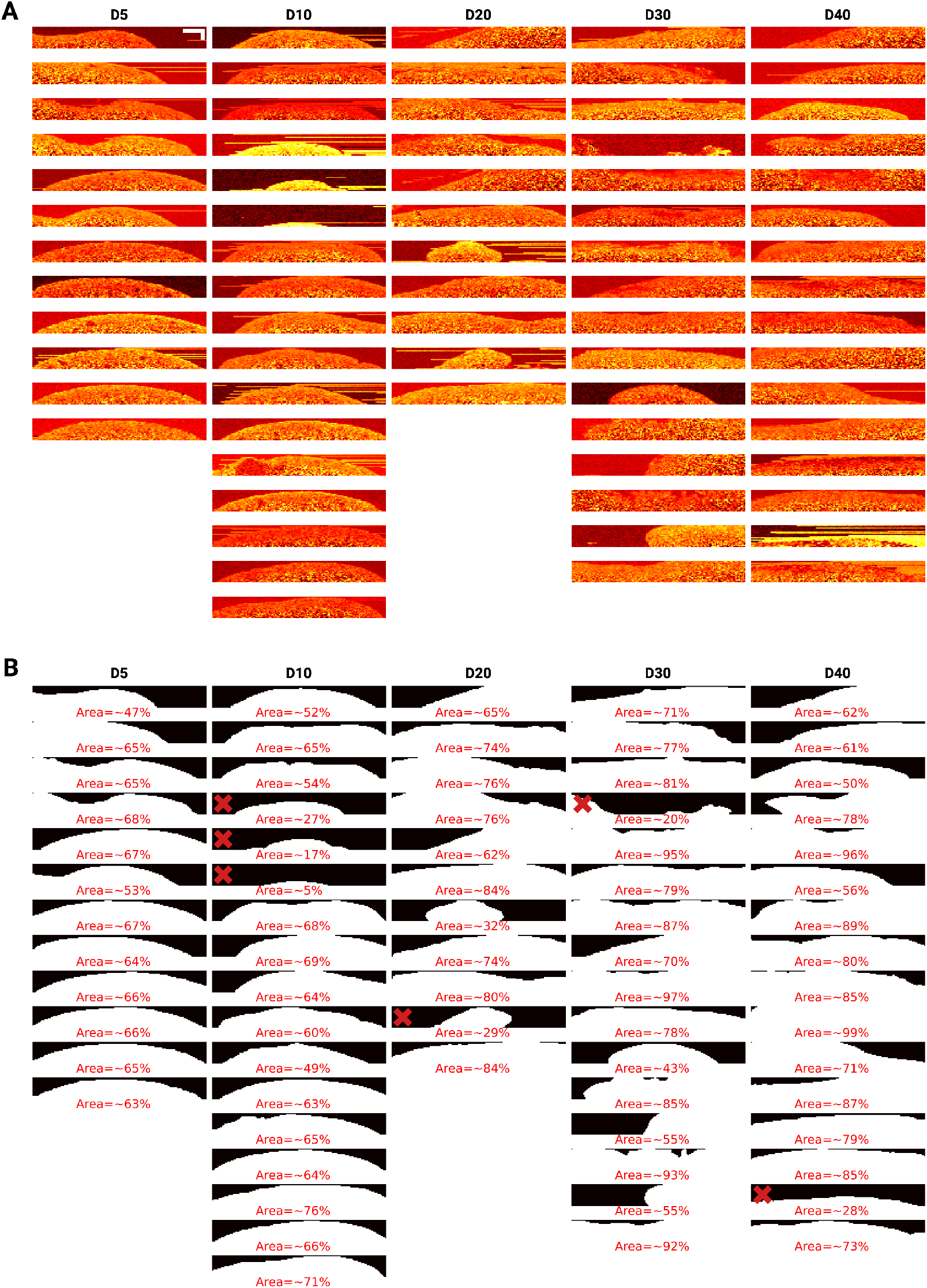
Preparation of timepoint Raman data for quantification analysis (as shown in Figures 5B–D in the main text). **A**, Intensity slices across the 1659 cm^−1^ band (linked to proteins and lipids) of the collected Raman imaging scans. Images were normalised for visualisation purposes. Scale bar: 100 µm (top left). **B**, Annotated organoid masks, along with estimated tissue portions. Scans with tissue areas smaller than 30% were not included in quantification analysis (marked with a red cross).

**Figure S15:**
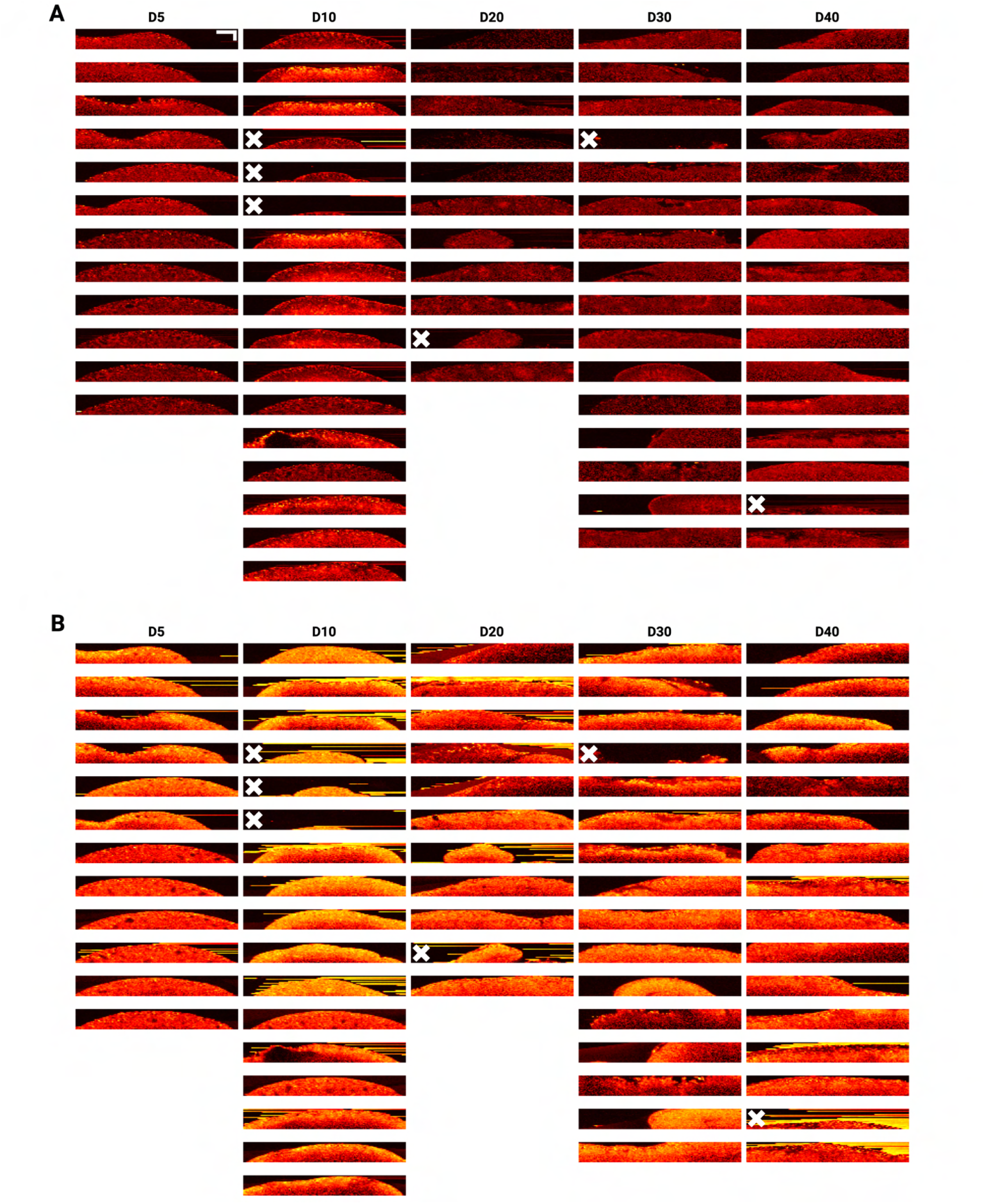
Abundance images of the two most abundant endmembers across timepoint RS scans (as shown in Figures 5B–D in the main text). **A**, Abundance images of endmember 1 (lipid-rich). Scale bar: 100 µm (top left). **B**, Abundance images of endmember 3 (protein- and nucleic acid-rich). Images were not normalised. Scans excluded from quantification analysis are marked with a white cross.

